# Multiple Loci for Foveolar Vision in Macaque Monkey

**DOI:** 10.1101/2024.02.01.578387

**Authors:** Meizhen Qian, Jianbao Wang, Yang Gao, Ming Chen, Yin Liu, Dengfeng Zhou, Haidong Lu, Xiaotong Zhang, Jiaming Hu, Anna Wang Roe

**Author notes:** Co-first authors: contributed equally to this work (Meizhen Qian, Jianbao Wang, Yang Gao). Corresponding Authors: Anna Wang Roe, Professor & Director, Interdisciplinary Institute of Neuroscience and Technology, School of Medicine, Zhejiang University, Hangzhou, China; Jiaming Hu, Assistant Professor, Interdisciplinary Institute of Neuroscience and Technology, School of Medicine, Zhejiang University, Hangzhou, China; Xiaotong Zhang, Professor, College of Electrical Engineering, Zhejiang University Hangzhou, China; Correspondence and requests for materials should be addressed to: Anna Wang Roe.

## Abstract

A common tenet of neural sensory representation is that species-specific behaviors are reflected in specialized brain organizations^1^. In humans and nonhuman primates, the central one degree of vision is processed by the foveola^2^, a retinal structure which comprises a high density of photoreceptors and is crucial for primate-specific high acuity vision, color vision, and gaze-directed visual attention^3,4,5^. In this study, we have developed high spatial resolution ultrahigh field 7T fMRI methods for functional mapping of foveolar visual cortex in awake monkeys. We provide evidence that, in the ventral pathway (V1-V4 and TEO), viewing of a central small spot elicits a ring of multiple (at least 8) foveolar representations per hemisphere. This ring surrounds a large area called the ‘foveolar core’. This is an area populated by millimeter-scale functional domains sensitive to fine stimuli and high spatial frequencies, consistent with foveolar visual acuity, as well as color and achromatic information, and motion. The unique position of the foveolar core suggests it may be a hub subserving higher order needs of foveolar function, such as integrating different spatial scales of representation, integrating local and global features in object perception, and bringing together the four quadrants of visual space. Thus, this elaborate re-representation of central vision signifies a cortical specialization for various foveation behaviors.

---

Across diverse species, the brain is known to contain representations of behavioral specializations^1^. In rodents the ‘barrel cortex’ is a specialization which processes tactile sensory inputs from each of the whiskers, providing essential information for object recognition and navigation^6^. The brain of a star-nosed mole has a specialized representation of the 22 tentacles on its nose used for actively touching and searching its environment for prey. Yet another example is the platypus whose brain contains specialized representations of the rows of electrical and vibrotactile receptors on its bill used to detect objects in the water.^7^ These cortical specializations thus anchor unique behavioral sensory activity to functional organizations in the brain.

Human and nonhuman primates also have characteristic specializations, but whether this is reflected in cortical organization is unknown. Here, to examine whether primate brains follow the principles observed in other species, we focus on a well-known and defining specialization for visual behavior in primates called the retinal *fovea* (center of gaze). Of particular interest is the *foveola*, the central most (central 1°) region of the fovea. This structure is characterized by an extremely high density (10 times the density of non-foveal retina) of photoreceptors needed for high acuity vision and color vision^3^. The foveola is also important for visually directed attention in primates, which is mediated by foveating (moving the eye) to objects of interest^8^. The essential role of central vision in common tasks such as reading and facial recognition predicts the existence of foveal brain circuits underlying learning and social behavior^9,10^. While there are extensive studies of extra-foveal brain areas, relatively little is known about foveal, and, in particular foveolar, brain organizations.

## Improving precision of MRI mapping

Early electrophysiological retinotopic mapping studies proposed a single locus where several foveal representations of early visual areas (V1, V2, V3, V4 and TEO) converge, termed a ‘foveal confluence’ ^11^. However, focusing on foveolar cortex has been challenging due to need for excellent fixation and fine precision mapping^16,17,13^. Electrophysiological studies in foveolar locations are difficult due to the extremely tiny (∼0.1°) receptive fields and ever-present small eye movements (micro-saccades). While eye movements can be largely removed in anesthetized animals, it is difficult to precisely locate (to within <0.5°) where on a screen the fovea is viewing. Thus, due to the precision required, basic questions remained unanswered or controversial. These included whether the foveolar representation is a single confluence or multiply represented across cortical areas^14,15,16^, what the foveolar cortical magnification factor is (a measure of the behavioral importance), and how distinct visual features such as color, shape, and motion are represented.

The advent of neuroimaging significantly advanced our understanding. The development of phase encoded fMRI mapping^17,18^ provided much insight into the layout of multiple visual areas in the ventral and dorsal visual pathways^19,20,21^, and revised the concept of foveolar confluence, revealing several foveolar loci along the anteroposterior extent of striate and extrastriate areas, both in humans^14^ ^15,20^ and in nonhuman primates^22,23^ (and, as was later shown, other confluences for the MT cluster^22^ and the posterior parietal cluster^24^). Many phase encoding fMRI studies illustrate retinotopic maps where vertical meridian (VM) and horizontal meridian (HM) apparently fall short of the center^25,26,27^, potentially indicating a need for further in depth study. In a very careful and systematic study, Schira^20^ and colleagues achieved this precision using fine phase encoding procedures designed specifically to examine foveolar representation in humans. They found that within the central 1°, there are smooth transitions along iso-eccentricities in the foveola and a clear phase discontinuity between dorsal and ventral V2 and V3 representations (also been borne out in monkeys studies^22^). Their maps also show the presence of some inhomogeneity in the very center (∼0.5°) of iso-eccentricity phase maps (**Extended Data Fig. 5A**). In another study in humans^16^, population receptive field mapping of foveolar cortex revealed similar findings, including the presence of inhomogeneity in the very central representation; interestingly, even with such high-resolution methods, the VM and HM ended short of the center (**Extended Data Fig. 5B**). These details suggest that the foveolar region may need further study.

Here, using fMRI and optical imaging methods, we have further examined the retinotopy and functional organization of foveolar cortex in Macaque monkeys. One of the primary challenges of studying foveolar representation is the high spatial resolution needed to answer these questions. We have thus turned to mapping of foveolar cortex in awake, behaving macaque monkeys trained to perform stable foveal fixation and studied in high field MRI. Following in the footsteps of others who have pioneered these methods^19,15,14^, we adapted previous methods. One of the novel approaches we took was to conduct our studies in a large bore size human ultrahigh field 7T MRI scanner; this provided sufficient space for a monkey to sit comfortably in a sphinx position and also allow for placement of instruments around the head. One of these instruments is a custom 16-channel surface coil^28^ that greatly enhances the signal-to-noise within a local cortical region, thereby permitting visualization of fine structures in the brain. An acquisition strategy with combined reduced-field of view imaging and high parallel imaging acceleration was used to maximize the spatial encoding capability of the MRI system. This is key to enabling acquisition of submillimeter resolution functional images (0.6-mm in-plane). To achieve the 0.1° precision needed to map the central visual 1°, we pursued two methods for mapping the foveolar cortex: (1) mapping with very fine foveolar stationary visual stimuli and (2) with phase encoding method, using fine expanding iso-eccentricity rings and rotating iso-polar stimuli.

## Stability of Acquisition

One major challenge of studying foveolar representation is minimizing eye movements. Two macaque monkeys were trained for several months to maintain stable performance with little body motion (**Fig. 1A**) and few saccades (**Fig. 1B, left**) during scanning. Only data in which fixations remained within a 1° radius window >85% of the time and center of mass of the eye trace remained within 0.5° radius of the fixation point were used (**Fig. 1B, right**). With sufficient trial numbers, consistent and reproducible fMRI results were generated across different sessions (**Fig. 1C**). These procedures made direct retinotopic and foveolar mapping possible (**Fig. 1D, 1E**).

**Fig. 1.**
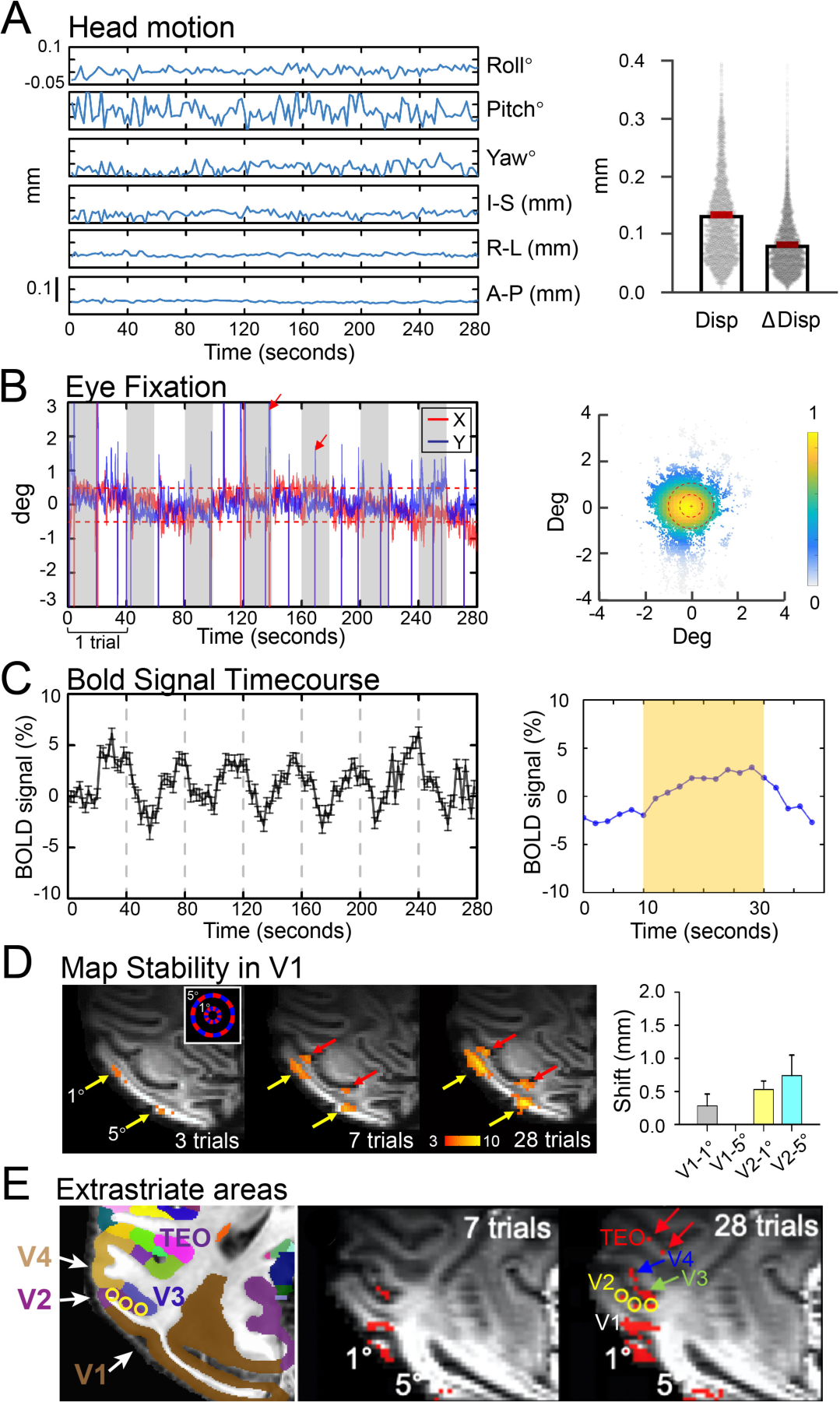
Acquisition Stability. (A) **Head motion**. *Left*: Motion in 6 dimensions. Roll, Pitch, Yaw, A-P (anterior-posterior), R-L (right-left), I-S (inferior-superior). *Right*: Head motion. Disp: Displacement relative to initial frame(mean = 120µm), ΔDisp: difference between two sequential frames (mean = 80µm). Averaged across 8 sessions. Monkey E. (B) **Eye fixation behavior.** *Left*. Eye position trace in 1 run. Red and blue lines: X and Y eye position coordinates, respectively. Two red arrows: blinks. Note eye blinks did not cause disruptions of the BOLD signal. *Right*. All eye positions in 4 runs (250hz, 4×280sec). Eye fixations were within 0.5 radius 94% of the time. Despite occasional saccades, across 4 runs (each trial 20 seconds blank and 20 seconds visual stimulus, 7 trials per run, total of 280 seconds), the center of mass of the eye trace was within 0.5° radius of the fixation point (right, central dotted red circle). With sufficient training, the monkeys maintained stable performance with little body motion and few saccades during scanning. (C) **BOLD signal timecourse**. *Left*: Timecourse of activated voxels during eye fixation in B. *Right*: Average BOLD timecourse. (D) **Map stability**. Maps obtained from 3, 7, and 28 trials. Left panel inset: visual stimulus: 1° and 5° eccentricity rings. Activation clusters remain stable across trials, indicating eye fixation is stable. Yellow arrows: V1. Red arrows: V2. Right panel: Shift in center of mass of activation clusters is minimal. (E) **Increased trial number reveals additional visual areas.** *Left*: Section from D99 atlas^51^. Activations in V1 (1° and 5°), V2 (yellow circles), V3 (green arrow), V4 (blue arrow), and TEO (red arrows). Same case as in D. Map threshholds: p<10^-3^. Error bars: Standard

## Foveolar Mapping

To determine the representation of the foveal region we first conducted visuotopic mapping (**Fig. 2)**. We identified the boundaries between cortical areas in the ventral pathway (areas V1, V2, V3, V4, and TEO). This was done for each monkey by mapping the representations of vertical (VM) and horizontal meridians (HM) using very narrow (0.15°) visual bars (**Fig. 2A, 2B**) as well as narrow (0.15°) iso-eccentricity arcs (**Fig. 2C, 2D**) during the fixation task (summarized for each monkey in **Fig. 2E, 2F**).

**Fig. 2.**
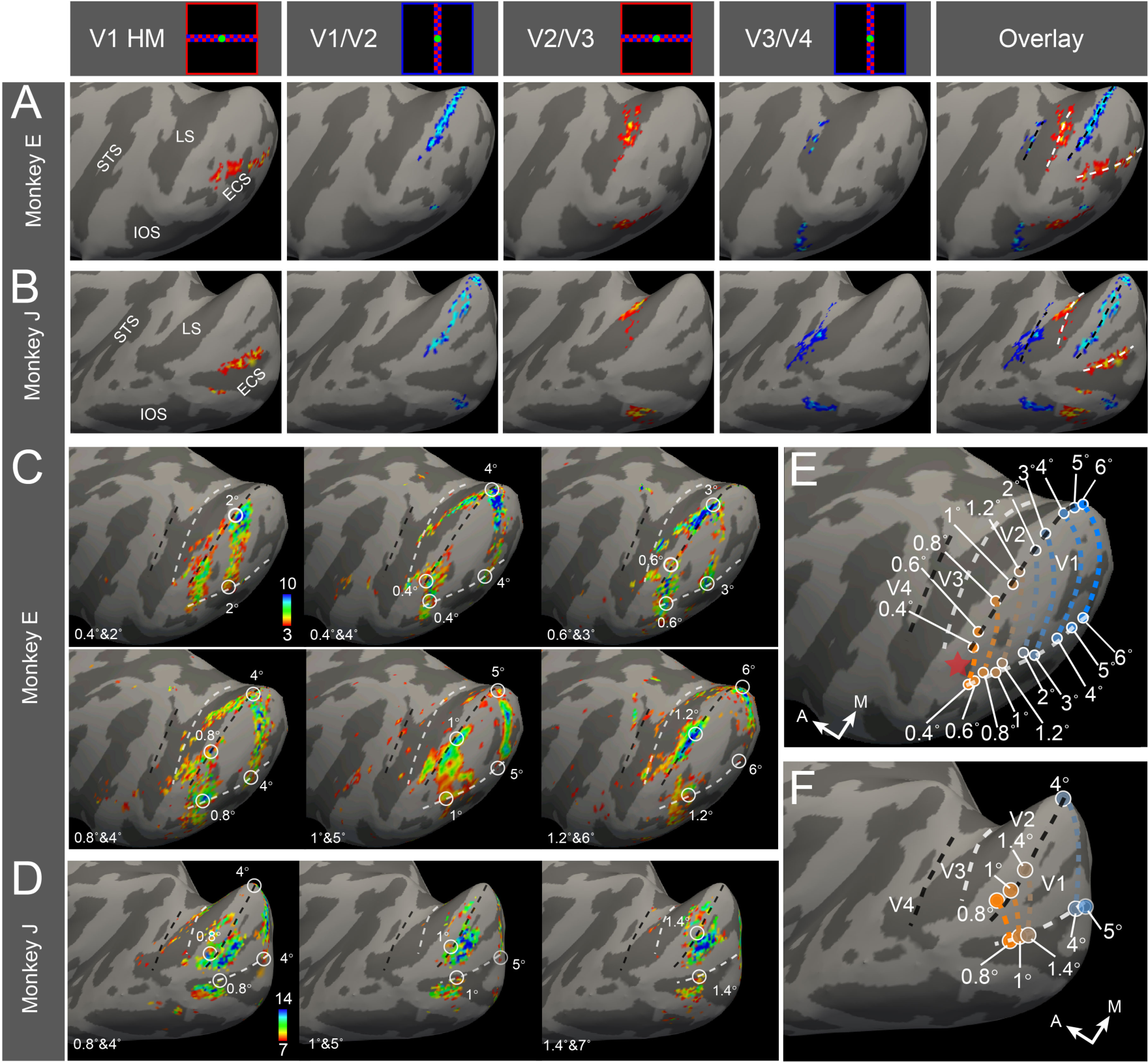
Mapping visual cortical borders by mapping HM and VM and iso-eccentricity mapping. (**A, B**) Top: Stimuli for imaging vertical meridian (VM) and horizontal meridian (HM) comprised horizontal and vertical 0.15° wide bands containing alternating saturated red-blue checkerboards (see insets at top). All images were acquired within the same sessions, but illustrated separately for each area (see Methods). For each monkey, the four panels at left were then overlain in rightmost images (Overlay). Black and white dotted lines: VM and HM, respectively. ECS: ectocalcarine sulcus; LS: lunate sulcus; IOS: inferior occipital sulcus; STS: superior temporal sulcus. Monkey E: p<10^-3^; Monkey J: p<10^-9^. (**C**) Iso-eccentricity maps from Monkey E. A paired stimulus paradigm (seven sets of paired rings: 0.4°&2°, 0.4°&4°, 0.6°&3°, 0.8°&4°, 1°&5°, 1.2°&6°) reduced the number of stimulus conditions needed by half and further confirmed maps were stable across sessions (e.g., similar 4° activations are obtained in 0°& 4°, 0.4°& 4°, and 0.8°& 4° maps). White circle: intersection location between iso-eccentricity and meridians. (**D**) Iso-eccentricity (0.8°&4°, 1°&5°, 1.4°&7°) maps from Monkey J. **E**&**F**. Summary of **C**&**D**.

To precisely identify foveolar locations, we used small stimuli placed at the fixation location. These comprised alternating saturated red/blue flashing spot stimuli (see Methods); to examine consistency of activation locations, three spot sizes were used (Monkey E: 0.4°, 0.6°, 0.8°; Monkey J: 0.6°, 0.8°, 1.0°). Given the distribution of eye positions during the fixation task, the most frequently activated locations are at the very center and thus the foveolar center should be at the *location of voxels with highest statistical significance*. Given the simple flashing visual spot stimuli used, the activation was strongest for early areas V1 and V2 and weaker at more anterior V3, V4, and TEO locations; the maps were examined using thresholds (3 p-values per area) appropriate for the activation levels of each cortical area. Below we describe how each foveolar location is identified in Monkey E (**Fig. 3**, Monkey J **Extended Data Fig. 1**). Division of voxels into V1/V2, V2/V3, V3/V4, V4/TEO, and TEO/FST regions were guided by locations of VM and HM (black and white dotted lines, respectively, shown in Overlay in **Fig. 3A**, last column) as well as slice views (**Extended Data Fig. 2)**. Below, we describe each foveolar location.

**Fig. 3.**
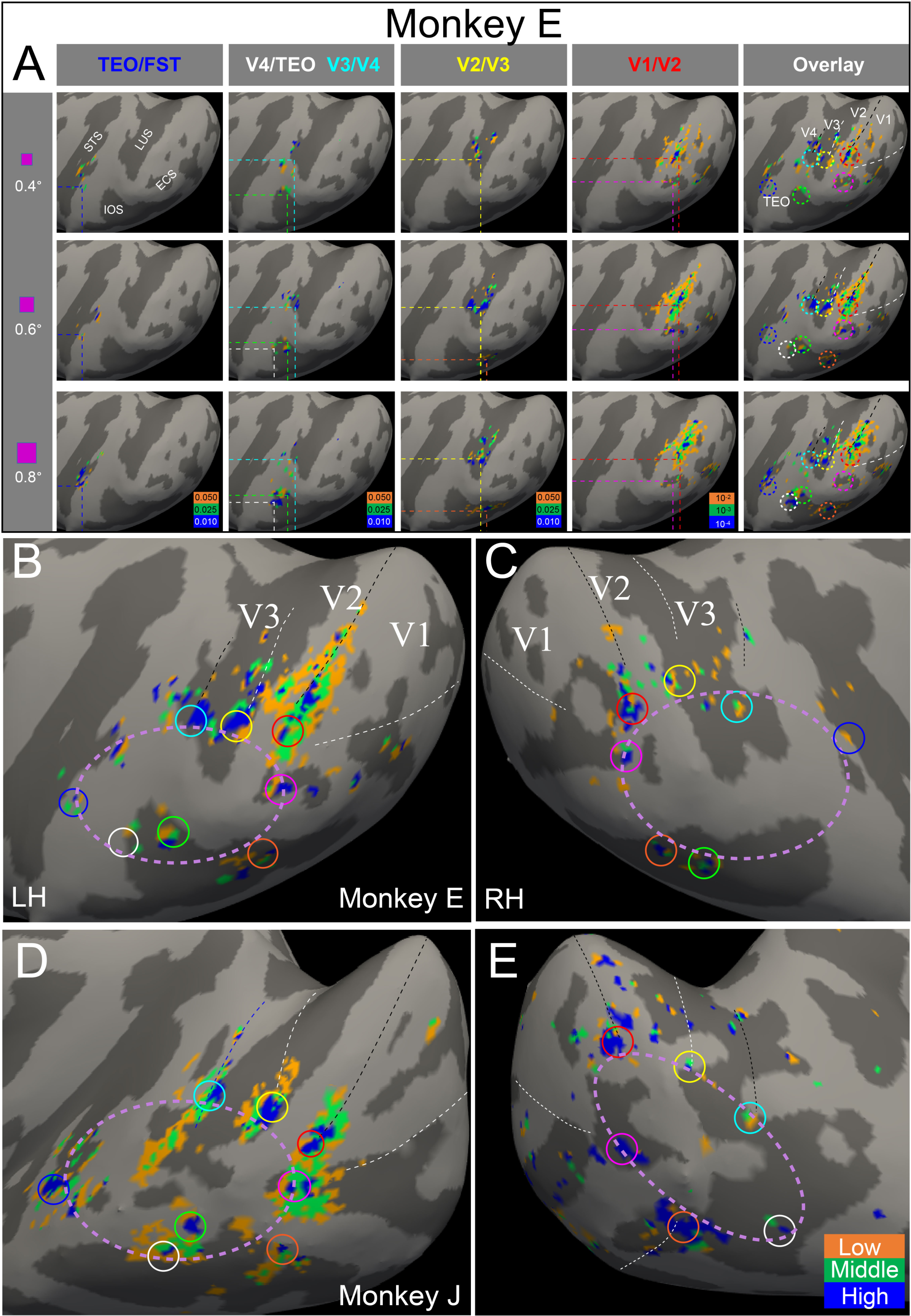
Determining locations of foveolar representation. (A) Activations to foveated small spot stimuli (0.4°, 0.6°, 0.8° flashing saturated red/blue squares alternating at 3Hz; shown in first, second, and third row, respectively, Monkey E). Shown in each column (TEO/FST, V4/TEO, V3/V4, V2/V3, and V1/V2) are the activation maps to each stimulus. Each panel: significant voxels at each of 3 thresholds [p-values indicated by color code: orange (lowest), green (middle), and blue (highest)]. For each visual area, the center of highest significance is consistent across spot sizes (dashed lines). Last column: the set of all foveola (colored dashed circles, overlay from 4 columns at left). Black and white dashed lines: VM and HM, respectively. Gyri: light gray; sulci: dark gray. ECS: ectocalcarine sulcus; LS: lunate sulcus; IOS: inferior occipital sulcus; STS: superior temporal sulcus. (B) Foveolar activations in Monkey E, right hemisphere (same as response to 0.6° in A). (C) Foveolar activations in Monkey E, left hemisphere. (D-E) Foveolar activations in Monkey J (D: left hemisphere, E: right hemisphere). See **Extended Data Fig. 1**. C-E: Small colored circles: foveolar activations in each cortical area. Lavender dotted oval: foveolar core.

### V1/V2 (fourth column)

In Monkey E, each of the 3 spot sizes activate a similar region (compare three rows in **Fig. 3A**). Examination of the activations at the lowest threshold (V1/V2 column, orange, 0.4° stimulus) reveals patchy activations across a centimeter-sized region; this activation region is anisotropic (more elongated along VM axis), consistent with previously observed anisotropies in V1 and V2 due to functional re-representations^29^ and/or to anisotropic representation along cardinal axes^30^. Although eye fixations away from the fixation location are relatively few (set by our criterion of <15% of fixations outside of 1° radius), we cannot exclude the possibility that they contribute to some of the activation seen at lower thresholds. At higher thresholds, quite focal activations are revealed across smaller regions of cortex (middle threshold, green: 3-8 mm; highest threshold, blue: 1-4 mm). We defined the “most foveal” locus as the lateralmost (since retinotopic center is located laterally on macaque visual cortex) activation at the highest threshold (blue) that was consistent across the 3 spot size maps. The dorsal locus overlay the V1/V2 border (red dotted lines) and another locus was slightly ventral to the HM in V1 (magenta dotted lines). Across the 0.4°, 0.6°, and 0.8° spot sizes, the locations are similar (see red dotted circles in Overlay) and are consistent with overall topographic (HM, VM) organization.

### V2/V3 (third column)

There were also two loci for V2/V3. The dorsal activation fell within the banks and depths of the lunate sulcus (LUS, yellow dotted lines) consistent with previous studies^31,32^, had a distributed patchy appearance, and was topographically consistent with a foveolar location along the V2/V3 border (see Overlay, dotted yellow circle). A second foveolar locus (evident for the 0.6° and 0.8° spots but below threshold for the 0.4° spot) fell in the inferior occipital sulcus (IOS) (cyan dotted lines, see Overlay dotted orange circle).

### V3/V4 (second column)

Two distinct foci are seen at the V3/V4 border, one on the anterior bank of the lunate sulcus (cyan dotted lines) and one at the tip of the IOS (green dotted lines). There are also a few patchy activations extending onto the cortical convexity (between the ends of the LUS and IOS and posterior to STS). The locations of these loci fall at the end of the V3/V4 border representing the VM (see Overlay, cyan dotted circle).

### V4/TEO (second column) and TEO/FST (first column)

TEO is a mid-tier area and is likely to exhibit somewhat weaker response to simple spots^33^. Despite this, a locus of patchy activation is visible at the tip of the IOS (white dotted lines, V4/TEO border) and at the posterior border of the STS (blue dotted lines, consistent with activation at the TEO/FST border^22^).

### Bilateral Representation

Since the fovea is known to be represented in both hemispheres, we also examined the foveolar representation in the contralateral hemisphere. The RF coil is typically placed slightly off center biased towards one hemisphere, so that we can capture as much of the ventral pathway as possible. The SNR on the contralateral hemisphere is therefore slightly lower, but significant BOLD signals can still be captured. In addition to the activation shown above (**Fig. 3B**, same as that shown in **Fig. 3A middle row**), foveolar activations in the contralateral hemisphere, obtained from the same sessions, are shown in **Fig. 3C** (see **Extended Data Fig. 2** for slices views). Each of the foveolar locations identified in **Fig. 3A** are indicated by circles of the corresponding color. Activation in TEO on the right side is present but weak (white circle in Monkey E). Similar data for Monkey J is shown in **Extended Data Fig. 1** and summarized in **Fig. 3D** and **3E**.

In sum, these methods reveal precise foveolar activation locations. Foveolar activation often comprised one to a few distinct patches. The locations of these activations were stable across multiple sizes of spot stimuli and were consistent across multiple thresholds. Due to the mapping precision, these data suggest that, in contrast to what was previously viewed as a singularity, there are at least 8 distinct nodes of foveolar representation, one near the end of each VM and HM for dorsal and ventral V1/V2, V2/V3, V3/V4, V4/TEO, and TEO/FST. This mapping is simultaneously observed in both hemispheres. Thus, the foveolar representation is not a single point or confluence, and is more than one per visual area, as previously suggested. Moreover, these findings suggest there is cortical territory bounded by these foveolar representations that is not within the visuotopically mapped regions (cf. **Fig. 8**). We provide a schematic ring (dotted lavender ring in **Figs. 3B-3E**) around this central region and refer to it as the *foveolar core* (or *core*). This ring is not meant to be a specific border but an indication of the region of cortex bounded by the foveolar loci.

We considered whether these findings could be an artifact of experimental procedures or data processing. (1) To eliminate the potential artifacts that the foveolar core was due to the subtraction of fixation cross, we examined single condition maps of spot and fixation cross (shown in **Extended Data Fig. 3)**. Single condition maps (no subtraction) to small foveal color **(Extended Data Fig. 3A)** and to the fixation cross **(Extended Data Fig. 3B)** evoke no activation in foveolar core (dotted pink oval, defined by multiple foveolar activations, yellow circles). Thus, the absence of signal in the core remains, and is not due to fixation cross subtraction. (2) We also provide evidence that the lack of activation in the core is not related to variability in fixation or to the possibility that this result is produced by the edges of the stimulus spot (**Extended Data Fig. 4**). With larger eye movements, the edges of the spot stimulus should activate more peripheral loci on the cortex. However, this is not observed. The activations on the cortex remain very similar across poorer (85% within 1°) and better (95% within 1°) eye fixation behavior Second, it predicts that a larger (e.g., 0.8°) would activate more peripheral locations on the cortex than a smaller (e.g., 0.4°) stimulus. However, as shown in **Fig. 3** and **Extended Data Fig. 1**, the foveolar loci are stable across 3 spots sizes, in each of 2 monkeys. (4) Blurring due to eye movements or other motion related artefact would only serve to make the activations broader and less separable, making our results an *overestimate* of the true foveolar activation size.

## Phase encoding reveals a nonhomogeneous region and distinct foveolar core responses

As phase encoding is the standard method for mapping retinotopy, we have also conducted phase encoding for the central 3° in a fixating monkey (Monkey J). Schira and colleagues^20^ studied the foveolar region in humans with phase encoding using very fine stimuli, high resolution functional imaging (1.2 mm isotropic), and sophisticated analysis. We attempted a similar study using a continuous sequence of iso-eccentric thin (0.15° wide) rings and narrow (5°) isopolar wedges. The resulting phase maps are shown in **Fig. 4A** (iso-eccentricity) and **Fig. 4B** (iso-polarity); the foveolar core is indicated by a dashed lavender ring and foveolar loci by colored circles (**Fig. 3**). Consistent with phase encoding studies^18,25,17,20,22^, in the iso-eccentricity map there is a systematic phase shift in V1, V2, V3 on the lateral operculum, with the central-most (least phase shifted) voxels (red) appearing most lateral on the cortex. This is further illustrated by the shifting time-courses across ROIs #1-6 along the V1/V2 border (**Fig. 4D, left column,** graphs #1-6, see shifting red arrowheads); the regularity of the time-courses and small degree of variability in these traces indicate that eye fixation was excellent.

**Fig. 4.**
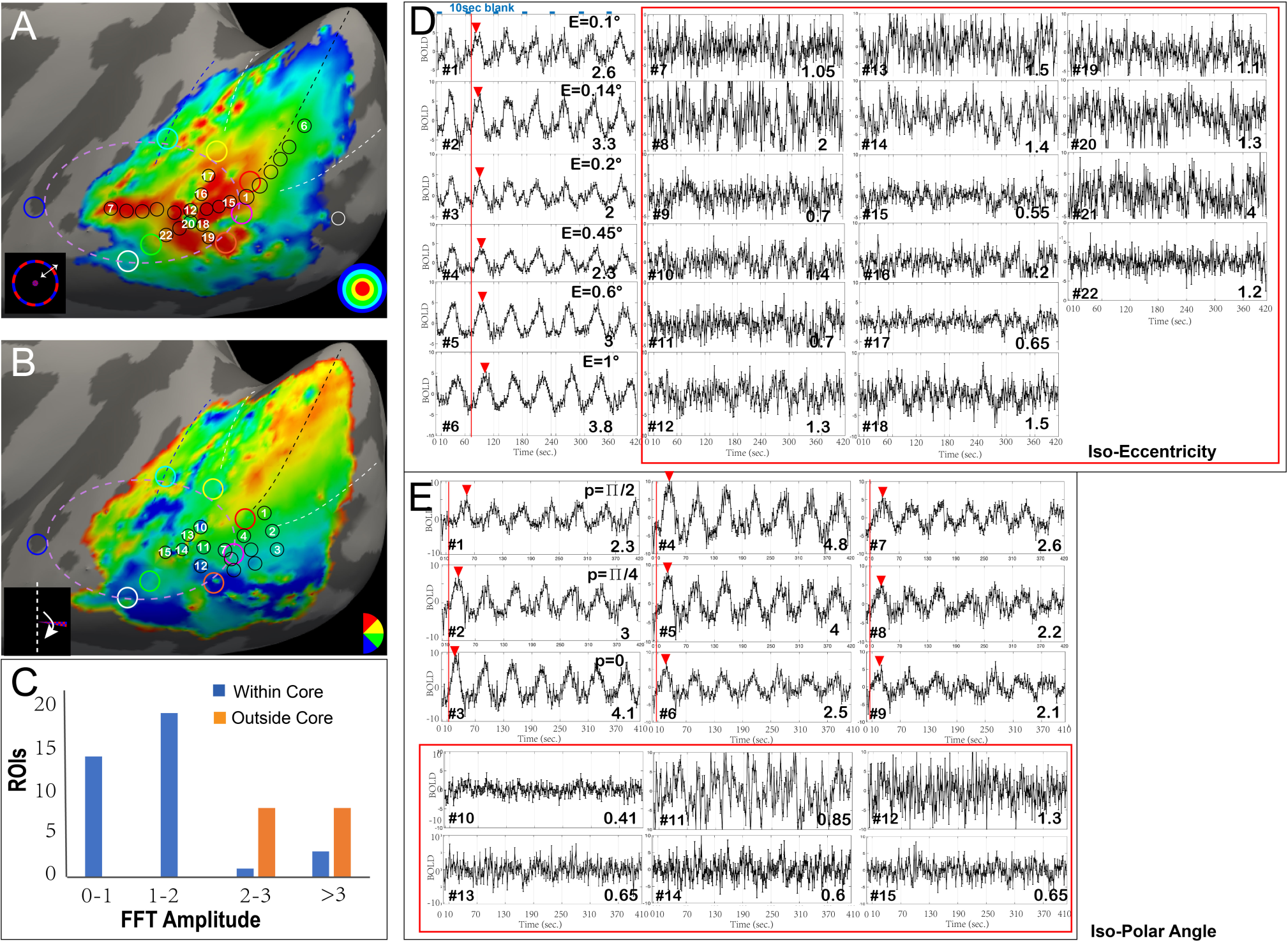
Phase encoding using fine iso-eccentricity and iso-polar stimuli in visuotopoic V1 and in foveolar core. **(A)** Iso-eccentricity map. Inset: Iso-eccentricity stimulus: continuous increasing size of iso-eccentric rings (each ring 0.15° wide; 1 trial: 50 rings over 3°, 0.06° per ring; 1 trial: 50 seconds, 10 second ISI; fixation point: 0.05° white dot which is constant, total 21 trials). **(B)** Iso-polar map. Inset: Iso-polar stimulus: continuous rotating checkerboard wedge (5° wedge, rotated over 50 sec from 90° to −90° in contralateral visual field, size 3°, total 21 trials. Colored circles: locations of voxel time-courses shown in C and D. **(C)** Histogram showing FFT amplitude at stimulus frequency is weaker in ‘inside core’ compared to ‘outside core’ samples. FFT amplitude inside core: mean=1.2; Outside core: mean=3.0. Chi-squared statistic, p< 0.01. **(D)** Iso-eccentricity time-courses (BOLD%). ROIs #1-6 in V1 along the VM. Red arrowheads (center of mass of peak) indicate shifting of peak as ROIs approach center. ROIs #7-15 in foveolar core along an anteroposterior axis through the core, ROIs #18#19 along a posteroinferior axis, ROIs #20-22 along a anteroinferior axis, ROIs#16#17 along a posterosuperior axis. **(E)** Iso-polar time-courses (BOLD%). ROIs #1-9 along 3 iso-ecc rings in V1, ROIs #10-12 in foveolar core parallel to iso-ecc rings in V1. Note that since dorsal cortex represents ventral fields, the phase shift due to the wedge approaching the VM in the ventral field will result in a shortening latency to response (red arrowheads). Red boxes: Time-courses from ROIs in the core region. Numbers in right corner: FFT amplitude indices. See **Extended Data Fig. 6**.

We hypothesized that if the foveolar center were within the core, we would observe continuously shifting phase from the several foveolar loci towards the center. We therefore sampled phase response at locations through the most central (red) regions of the foveolar core starting from (1) the dorsal V1/V2 foveolar locus (ROIs #7-15), (2) the dorsal V2/V3 to the ventral V2/V3 foveolar loci (ROIs #16-19), and (3) the ventral V3/V4 foveolar locus (ROIs #20-22). [Note anterior V4/TEO locations were outside our imaging FOV.] As shown in **Fig. 4D right three columns**, we found that these samples exhibited surprisingly noisy responses which are poorly phase-locked to the stimulus (**Fig. 4D**, see FFT amplitude indices **Extended Data Fig. 6**).

In the iso-polar map (**Fig. 4E**), outside the foveolar ring the ROIs were well modulated and showed phase shifting (**Fig. 4E**, top 3 columns ROIs #1-3, #4-6, #7-9, see shifting red arrowheads). Again, inside the core region, sampled from a parallel arc just inside the border of the core region (**Fig. 4E**, ROIs #10-12) and in the central zone of the core region (**Fig. 4E**, ROIs #13-15), the traces were noisy and not well modulated. This is supported by significant differences in the peak amplitude of the stimuli frequency of ‘Within Core’ and ‘Outside Core’ samples (**Fig. 4C, Extended Data Fig.6**). Thus, whereas our methods could detect phase shifting outside the core, the responses in the core appeared noisy and poorly phase-locked. If the noisy signals were due to eye movements, the foveolar traces from ‘outside core’ samples would also be noisy, inconsistent with the stable traces obtained. It may be possible that neurons in the core prefer even smaller stimuli (<0.15°, see also **Fig. 5**); however, the ‘within core’ traces do exhibit responsiveness to the stimuli (large magnitude fluctuations), but have poor phase correlation. While the nature of the noisy traces remains to be examined, these data suggest that responses in the core region are qualitatively distinct from those in retinotopic regions of visual cortex.

**Fig. 5.**
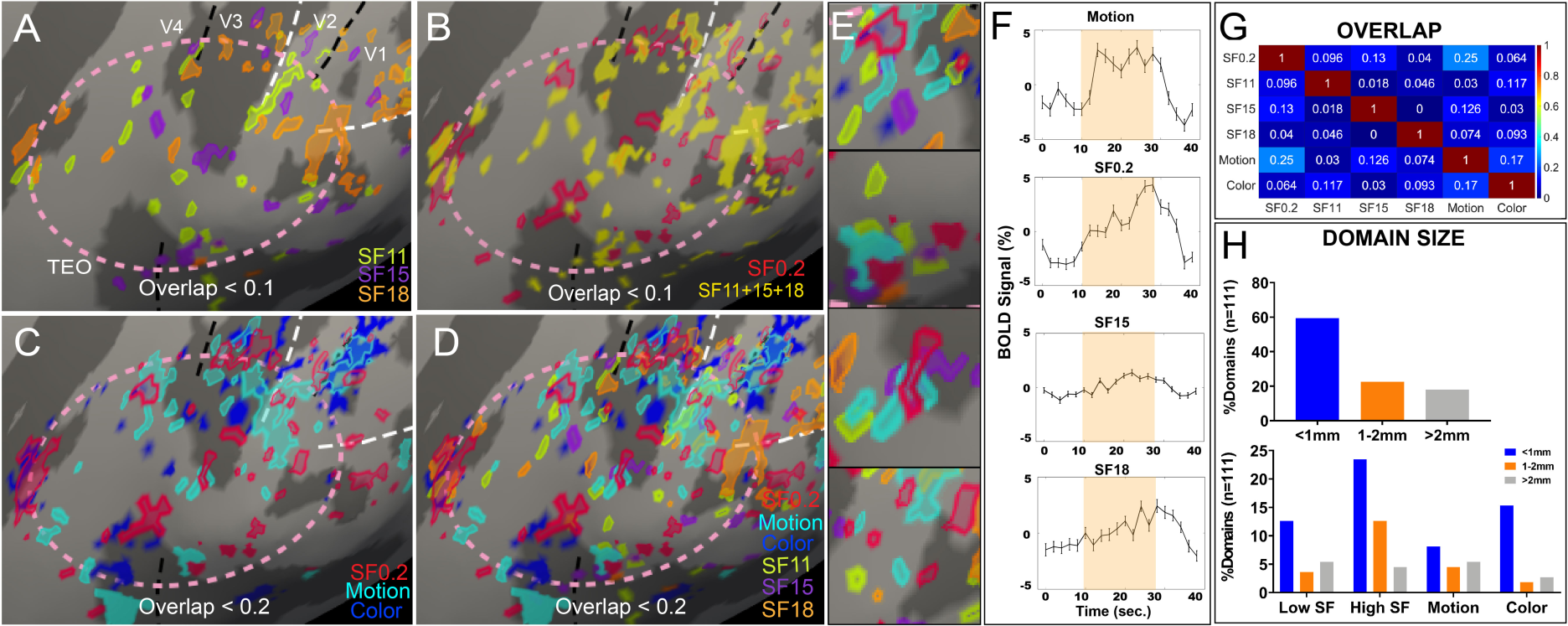
Mesoscale functional domains within the foveolar core. Color-coded responses to different visual stimuli. (**A**) Activations to achromatic high spatial frequency gratings (SF, in cycles/deg). Yellow: SF 11. Purple: SF 15. Orange: SF 18. (**B**) Overlay of all high SF (yellow: SF11+SF15+SF18) and one low SF (red: SF 0.2) activations. (**C**) Overlay of low SF (red: SF 0.2), motion (cyan: clockwise moving dots), and color (blue: 0.8° flashing spot) activations. (**D**) Overlay of all domains. (**E**) Four fields of view from (**D**) showing multiple types of functional domains within a small region. (**F**) Averaged time-course of each domain type selected from 3-5 clusters within the core. Error bar: Standard error. (**G**) Overlay indices between different populations of domains show little overlap between domains (overlap indices 0.09, range 0-0.25). (**H**) Domain size distribution (total number: 111, see Methods, **Extended Data Fig. 8**).

We also observe a nonhomogeneous variation (reds to greens) in the core, consistent with previous studies in foveal regions in humans^20,14^ In fact, close perusal of these studies reveals the presence of similar foveolar loci along a ring surrounding a central nonhomogeneous region (see **Extended Data Fig. 5)**. Although statistical thresholding was used, it would be important to see the strength of correlation of phase encoding for the central field voxels (cf. **Extended Data Fig. 6**). We also found potentially similar published results using retinotopic mapping by pRF analysis^14^. The central area in these maps was not homogeneous and exhibited focal red loci which fell at the end of the VM and HMs and on a ring that encircled what may be the foveolar core (**Extended Data Fig. 5B**). Other published phase encoded maps also illustrate VM and HMs that stop short of a common center^14,26^. We suggest that the core region does respond to foveolar stimuli, but their receptive fields may not be topographically ‘foveola only’ (e.g. the lack of response to small focal stimuli and, in some cases, preference for large stimuli, see **Fig. 4** on functional domains).

## Are there functional domains within foveolar cortex?

One of the defining features of visual cortical areas is their functional organization for different parameters of visual information. Well known are the stripes of V2 which range in size from 1-2 mm in width and the distinct sets of feature specific submillimeter-sized domains within the thin, pale, and thick stripes^29,31,34^. While, imaging millimeter-or submillimeter-scale modular structures with fMRI in monkeys has been challenging^31,23^, we illustrate here that V2 color (thin) stripes are well mapped by color vs achromatic gratings, but fail to evoke much activation in the foveolar core (**Extended Data Fig. 7**). We then proceeded to explore the foveolar core response using a small battery of visual stimuli, including high and low spatial frequencies, simple color and achromatic stimuli, and motion dot stimuli.

### High spatial frequency domains

A defining feature of foveolar function is its high visual acuity, marked by neuronal responses to high spatial frequencies visible only to foveolar vision^35^. Neurons within foveolar locations of macaque V1 have extremely small receptive field sizes (predominantly 10-20’ in size^36^) and exhibit peak spatial frequency tunings up to ∼5-8 cyc/deg^37,38^. Peak spatial frequency tunings based on fMRI BOLD signals reach ∼3 cyc/deg^39^. However, perceptual detection and discrimination can reach SFs well over 10 cyc/deg in humans^35^. Thus, a potential function of the core region may be to integrate lower order SFs to achieve high spatial acuity^35,40^. In this section, we use the term ‘central’ to mean foveolar, not in the topographic sense, but in the functional sense of central visual perception. To capture central spatial frequency responses, we examined response to high spatial frequency (SF) achromatic gratings (SF11, SF15, and SF18 cyc/deg), and, for comparison, to low spatial frequency achromatic gratings (SF0.2 cyc/deg) and to small flashing color spots. Monkeys maintained excellent fixation behavior during presentation of these stimuli. As shown in **Fig. 5A**, activations to SF11 (yellow-green), SF15 (purple), and SF18 (orange) gratings revealed patchy activations scattered throughout the core (dashed oval). Moreover, SF11, SF15, and SF18 showed little overlap, suggesting they are distinct functional domains (**Fig. 5A**, compare yellow-green, purple, and orange; overlap index for all comparisons < 0.05, **Fig. 5G**). In sum, these high spatial frequency stimuli appear to effectively activate the region within the core region and exhibit distinct spatial activations.

We then compared these maps to those obtained with low SF0.2 achromatic grating, a spatial frequency typically used for studying surface properties like color and luminance. The SF0.2 responsive domains (**Fig. 5B**, red) were largely non-overlapping with the high spatial frequency domains (**Fig. 5B**, yellow: sum of SF11, SF15, and SF18, overlap indices < 0.1). Achromatic SF0.2 domains (**Fig. 5C**, red) also had low overlap with domains activated by a small (0.8°) color stimulus (**Fig. 5C**, blue; overlap index 0.12). For comparison, response to small motion dot stimuli (2-degree patch of dots rotating clockwise) (**Fig. 5C**, cyan) elicited activations with little overlap with SF0.2 achromatic (red, overlap index = 0.04) and low, but slightly larger, overlap with color domains (blue, overlap index = 0.16). Domains of different types appeared within small regions within close proximity to one another and did not appear to coalesce into stripes or bands (a few examples shown in **Fig. 5E**). Averaged time-courses of different domains within the foveolar core exhibited reasonable BOLD time-courses (**Fig. 5F**), indicating these activations are not artifactual.

### Domain sizes

We also estimated the sizes of the activation domains. The area of each activation was calculated by measuring the number of nodes within the activation when overlaid onto the brain surface mesh^41^ (see **Extended Data Fig. 8**). Given the 0.6 mm in-plane resolution, we conservatively classified domains as <1 mm, 1-2 mm, and >2 mm in size. Domain diameters (the longest axis) revealed that they are largely < 2 mm in size (**Fig. 5H, top,** 82% ≤ 2 mm in diameter) and holds for different functional domain types (**Fig. 5H**, **bottom**).

Thus, remarkably, what originally appeared as a region of cortex with little response to foveolar spot stimuli, is actually a region densely populated with spatially distinct functional “domains” of different feature selectivity (**Fig. 5D** shows all domains overlain, all overlap indices < 0.15). Unlike the stripe and band organizations in early visual areas, each of the domain types appear to be distributed broadly across this central zone, resulting in a mosaic-like functional architecture. We speculate that a larger stimulus set may fill more of the territory within the foveolar core.

## A view from optical imaging

To further examine the topographic and functional organization of the foveolar core, we conducted intrinsic signal optical imaging of this area in anesthetized monkeys. To functionally localize the foveal representation in anesthetized monkeys, determination of the foveolar location was determined by sequential, iterative imaging of large interval to small interval vertical and horizontal bar stimuli to zero in on the precise x, y coordinates on the monitor (see Methods). As shown in **Fig. 6E**, the lateral visual cortex was exposed, including lateral parts (central 0.5°) of V1, V2, and V4, and part of TEO. Maps for ocular dominance revealed the V1/V2 border (**Fig. 6A**, fine dotted line). Mapping orientation preference (e.g., horizontal vs vertical) revealed orientation maps in V1, orientation stripes in V2 [corresponding to the thick (‘disparity’) and pale (‘higher order orientation’) stripes, regions between the yellow arrows], as well as larger orientation (‘curvature’) responsive bands in V4 (**Fig. 6B**, outlined in green). Color vs achromatic response revealed blobs (‘color’) in V1 (**Fig. 6C**, pattern of dark dots), thin (‘hue and brightness’) stripes in V2 (yellow arrows), and larger bands of ‘contextual color and brightness’ response in V4 (outlined in red) (**Fig. 6C**). The color responsive stripes in V2 aligned well with dark cytochrome oxidase histology (**Fig. 6D**, 3 stripes indicated by yellow arrows). Each of areas V1, V2, and V4 is characterized by distinct and well documented functional organizations^42,43,44^. These establish the quality of our optical imaging methods.

**Fig. 6.**
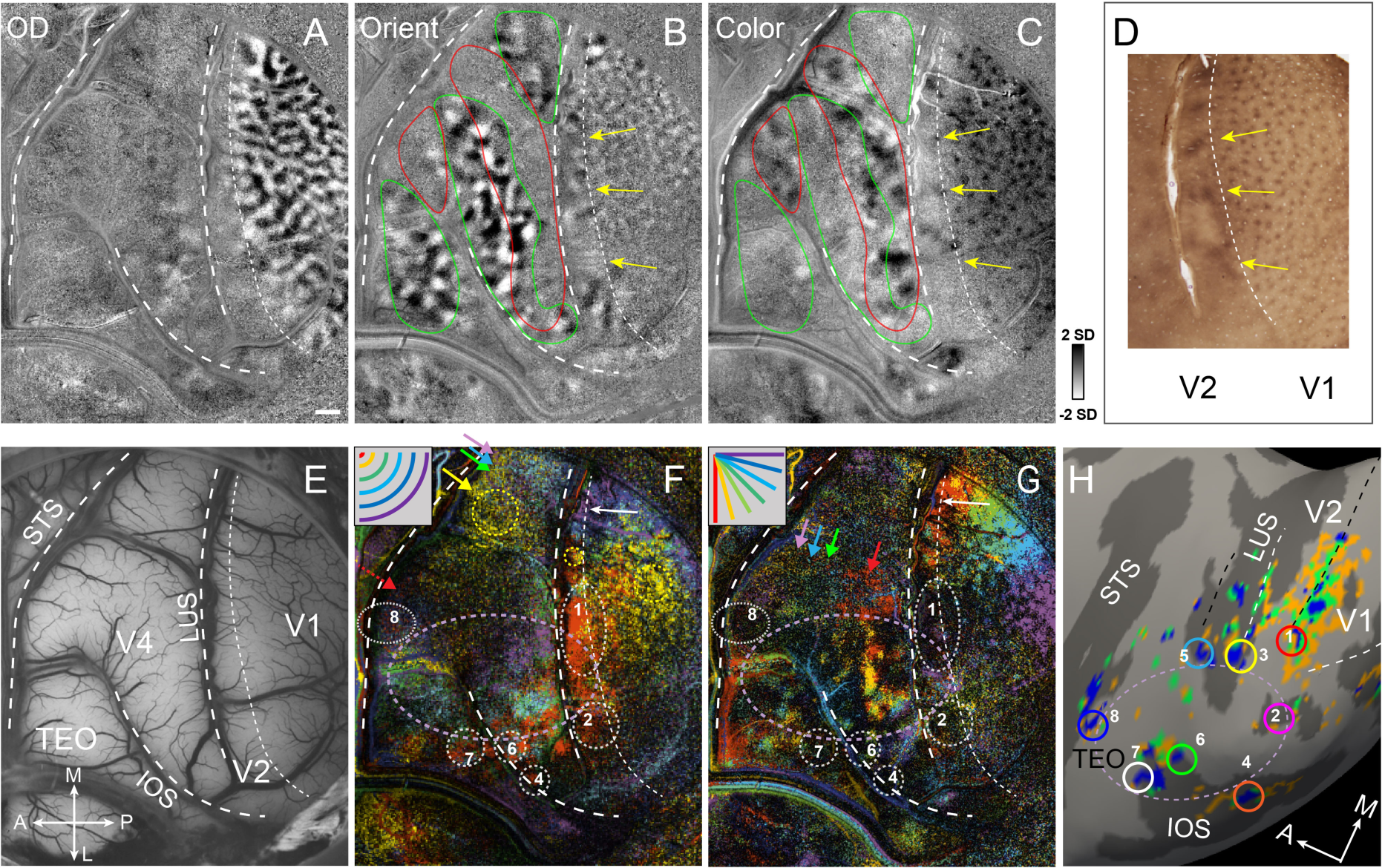
Multiple foveolar representations revealed by OI. (**A**) Ocular dominance (OD) map. Scalebar = 1 mm. (**B**) Orientation map (45° vs. 135°). Green circle: V4 orientation band. Red circle: V4 color band. (**C**) Color map (color vs achromatic). Yellow arrows: color stripes in V2, same locations as stripes shown in D. (**D**) Cytochrome oxidase stained stripes in V2 align well with color stripes in C (yellow arrows). (**E**) Blood vessel map over visual areas V1, V2, V4, and TEO. M: medial, L: lateral, A: anterior, P: posterior, LUS: lunate sulcus, STS: superior temporal sulcus, IOS: inferior occipital sulcus. (**F**) Color-coded eccentricity map (see inset: red 0.01°, yellow 0.10°, green 0.20°, cyan 0.30°, blue 0.40°, purple 0.50°. Lavender dashed circle: foveolar core. Colored arrows: shifted eccentricities. (**G**) Color-coded iso-polar map (see inset: red 0°, yellow 15°, light green 30°, dark green 45°, cyan 60°, blue 75°, purple 90°). Numbered white dotted circles: foveolar locations at V1/V2 (#1 and #2), V2v/V3v (#4), V3d/V4d (#5), V3v/V4v (#6), V4v/TEO (#7), and TEO/FST (#8). Location V2d/V3d (3) is within the lunate sulcus and not visible in OI. Colored arrows: shifted polar angles. (**H**) 7T MRI of all foveola in Monkey E (left hemisphere, from Fig. 3B) numbered with corresponding location in **G**. Orientation of all OI maps were rotated to correspond with MRI results in shown in H.

*Central 0.5° visuotopy (**Fig. 6F**)*. To determine foveolar representations, we conducted visuotopic mapping using thin (2 pixel wide) (1) iso-eccentricity arcs at 0.01°, 0.1°, 0.2°, 0.3°, 0.4°, 0.5° eccentricities (**Fig. 6F** inset, red, yellow, green, cyan, blue, purple, respectively) and (2) iso-polar lines, from vertical meridian to horizontal meridian (90°, 75°, 60°, 45°, 30°, 15°, 0°) iso-polarities (**Fig. 6G** inset, red, yellow, light green, dark green, cyan, blue, purple, respectively). These generated vector maps (color of each pixel reflects preference for one of the respective iso-eccentric and iso-polar conditions; for single condition maps see **Extended Data Fig. 9**). In V1, iso-eccentricity of the central 0.5° maps mediolaterally (**Fig. 6F**, from purple to blue/green, to yellow to red) and iso-polarity roughly posterior-to-anterior (**Fig. 6G**, from purple to blue to green to yellow to red). V2 (which only has 2 mm wide region visible on surface posterior to lunate sulcus LUS) exhibits a similar mediolateral mapping of iso-eccentricity (**Fig. 6F**), and shares the VM representation at the V1/V2 border (**Fig. 6G**, white arrow). The yellow dashed circles in F indicate corresponding locations of 0.1° eccentricity in V1, V2, and V4. Note that in the iso-polar color code, the foveal center responds to all the iso-polar conditions; this leads to no preference for any single condition and therefore results in black-colored pixels. Thus, the corresponding foveolar representations (foveolar loci #1 V1/V2d and #2 V1/V2v) appear red in the iso-eccentricity map and black in the iso-polarity map.

In V4, where receptive fields are larger than in V1 and V2, the map is weaker in response to the fine iso-eccentricity and iso-polar stimuli, but still consistent with known V4 topography. The iso-eccentricity topography runs mediolaterally (in **Fig. 6F**, most medial are purple, blues, and greens, with yellow pixels further lateral, colored arrows). Even more lateral is a large dark region (likely indicating poor response to central 0.01° stimulus) consistent with the foveal V4 location (dashed red arrow). In the iso-polar map, V4 exhibits weak responses, but an anteroposterior iso-polar topography can be discerned (in **Fig. 6G**, from purple to blue/green to red, colored arrows). Within TEO, the topography is less clear, and also weak to central stimuli.

### Foveolar centers and foveolar core

Two foveal centers at the V1/V2 border (**#1 V1/V2d** and **#2 V1/V2v**) are seen as red in **Fig. 6F** and black in **Fig. 6G**. Foveolar loci **#4 (V2/V3v**, at dorsal lip of IOS), **#6 (V3/V4v**, more anterior on dorsal lip of IOS) and **#7 (V4/TEOv**, anterior on ventral lip of IOS) also appear as red in **Fig. 6F** and black in **Fig. 6G**, And **#8 (V4/TEOd**) is visible on the surface and may extend into the STS (fewer red pixels in Fig. 6F due to weak response to tiny foveal stimuli). Foveolar loci **#3 (V2/V3d)** and **#5 (V3/V4d)** are not visible as they are within the lunate sulcus. The locations of #6, #7 and #8 are consistent in location with the corresponding loci shown in fMRI. In the core region (approximated by the lavender ring), there are intermingled patches of yellow, green, blue, red, purple, as well as dark regions.

In sum, the primary finding is that the optical imaging data largely is consistent with the fMRI findings: (1) there are multiple foveolar loci (dark circled loci in **Fig. 6G**), (2) the visuotopic maps, most clear in V1 and V4, do not continue into the core region, and (3) there are patchy intermingled activations in the core region. Although these data are from different animals in the fMRI data, these foveal centers provide support for those identified via fMRI (see corresponding numbers in **Fig. 6H**). Following this reasoning, the lavender dashed oval region in the optical image in **Fig. 6F** and **6G** indicates the approximate location of the foveolar core. Due to the folding of the cortex, the region corresponding to the foveolar core in the optical image may appear smaller than in fMRI. Thus, with a completely different imaging methodology with different spatial resolution, similar results are obtained, and further strengthens these findings.

## Foveolar Cortical Magnification

This finding introduces new questions regarding how much cortex is devoted to foveolar representation. Cortical magnification factors (CMF, defined as millimeters of cortical area devoted to processing a degree of visual angle) for each area has been well studied^45,46,47,48^; however, detailed data from the central 1 degree remains lacking^49^. CMFs calculated from retinotopic mapping (**Fig. 2E and 2F**) are shown from 2 monkeys in **Fig. 7A** and **7B**; these provided direct and precise fMRI measurements of cortical magnifications for the central 1°. CMF values at the very center reached up to around 24.6 mm/deg (red boxes), extending values from previous studies outside the central 1 degree (colored lines^46,49,50^). Differences between the CMFs of V1 in the 2 monkeys are consistent with previously reported inter-individual variability^43^ (previous values ranged from 4.5 mm/deg^51^ to 30 mm/deg^36^). From optical imaging data (**Fig. 6**), calculation of the CMF in V1 in the central 0.5° reaches about 16mm/deg (calculated from **Fig. 7C**). Further calculations from the OI data (see **Fig. 7F**), the area of the central 0.1° (yellow pixels) is ∼15mm^2^, resulting in an areal CMF of 1500 mm^2^/1°, and of the central 0.01° (for V1 half of the red pixels in V1/V2d) an area of ∼2.2 mm^2^, resulting in 22000 mm^2^/1°. The greater precision of optical imaging has provided better resolution for evaluating the relative increase in cortical magnification, illustrating that, rather than leveling off, the CMF is much higher at the most central visuotopic locations. Estimating area, measured from fMRI (see Methods) a single quadrant of the central 1° occupies an area of ∼70 mm^2^ in V1 or ∼140 mm^2^ for a single hemisphere of V1. If we take a very crude approximation that V2, V3, V4 have a similar area of representation (V2 and V3 may have slightly greater CMF^20^), then that number becomes 4 times larger or 560 mm^2^.

**Fig. 7.**
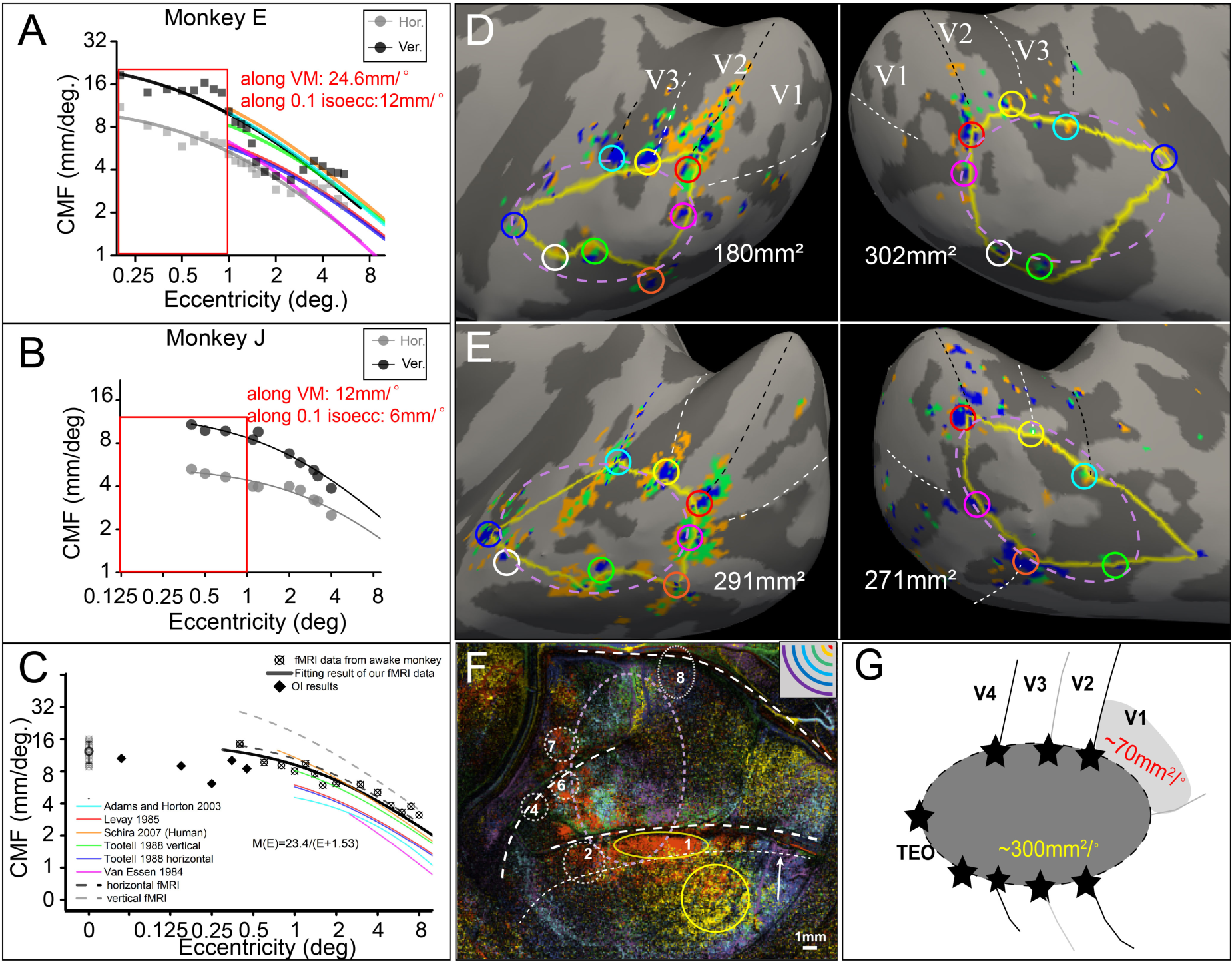
Foveolar core has large cortical magnification (CMF). (**A-B**) *Linear CMF* in central 1° of V1 as a function of eccentricity for (**A**) Monkey E (above) and (**B**) Monkey J (below). From **Extended Data Fig. 1**. Colored lines from^14^. And (**C**) *Linear CMF* in central 0.5° of V1, from optical imaging data in **Extended Data Fig. 9**. (**D-E**) Foveolar core bounded by 8 foveolar loci. Yellow outlines: boundaries for area size calculation, approximated by pink dotted oval. (**D**) Monkey E. (**E**) Monkey J. (**F**) Optical image of central 0.01° and 0.1° (yellow circles). The CMF of the central 0.1° (yellow pixels) is 1500 mm^2^/1°, and of the central 0.01° (half of the red pixels in V1/V2d) is 22000 mm^2^/1°. (**G**) Summary schematic of topographic (light gray) and non-topographic (dark gray) foveolar representation. Numbers: estimated area (mm^2^). See text.

## Size of foveolar core

We were surprised that the HM, VM and iso-eccentricity stimuli failed to activate the foveolar core. The HM and VM which pass through the central-most point of vision did not activate this zone, suggesting these stimuli, under conditions of fixation, were not effective for the most central region. To estimate the area in the foveolar core, we connected the 8 foveolar loci identified in **Fig. 2** and calculated the circumscribed area (**Fig. 7D** and **7E**, yellow outlines). The approximate area for this zone is 180 mm^2^ and 302 mm^2^ (for the left and right hemispheres of Monkey E) and 291 mm^2^ and 271 mm^2^ (for that of Monkey J); these lead to rough estimates of the radius of the core regions of about 8-10 mm. To further check that our numbers are consistent with our optical imaging, the measured distance between the dorsal and ventral V1/V2 foveolar locations (#1 and #2 in **Fig. 5G**) is 7.2 mm (see **Extended Data Fig. 9H**). This is comparable to the surface distance between the two V1/V2 foveolar locations from the fMRI map (**Fig. 6H**; #1-#2: 7.2 mm; other example distances are #5-#6: 5.5 mm, #7-#8: 11 mm, #5-#7: 6.3 mm, #5-#8: 14.7 mm). Thus, both fMRI and optical imaging data provide areal estimates of hundreds of mm^2^. Note that the central-most stimulus of 0.01° (red pixels in **Fig. 6F** and **Extended Data Fig. 9H**) covers a large area extending over 7 mm along the V1/V2 border, resulting in a central-most linear cortical magnification of ∼700 mm/deg (**Fig. 7E**).

In sum, the foveolar core region is large (roughly 200-300 mm^2^ in size). Adding in the approximated area of foveolar V1-V4 of 560 mm^2^, the total area devoted to foveolar vision is then upwards of 800 mm^2^ (**Fig. 7G**, dorsal foveolar V1 shown in light gray). As a point of comparison, the area of V1^52^ in one hemisphere ranges from 800 mm^2^ – 1500 mm^2^.

## Discussion

Following on studies using precision mapping of foveolar visual cortex in humans^45,20,15,18^, we developed ultrahigh field fMRI imaging methods and use of small foveolar stimuli in well-trained fixating monkeys. To our knowledge, this is one of the few awake monkey studies at 7T^53,54,55^ and the second study obtained in a human 7T scanner^53^.

### Foveolar maps

Previous visuotopic mapping studies have characterized the central-most foveal region as a single foveal confluence^11,22^ (**Fig. 8A, upper panel**) or several foveolar loci arranged anteroposteriorly^20,48,56, 53,57,58,59,60,61,62^ (**Fig. 8A, lower panel**). Our revised view (**Fig. 8B**, red stars) suggests (1) there are *two* loci per area, one each for dorsal and ventral representations, resulting in 8 distinct foveolar loci per hemisphere (8 red stars). These 8 loci do not converge, are spatially distinct, and surround a relatively large cortical region (roughly 200-300 mm^2^ in size). We have considered the possibility that these results are due to vascular artifacts, imaging artifacts, eye movement related blurring issues, stimulation by edges of the stimuli, and the subtraction of fixation cross, but found these possibilities to be inconsistent with the data. We also mapped the central 1° using phase encoding in response to fine stimuli, which revealed systematic iso-eccentricity and iso-polar maps for visuotopic foveolar cortex, but, noisy responses with poor correlation to the phase of visuotopic stimuli in the foveolar core. This difference is not easily explained by eye movements as both foveolar loci and foveolar core responses were simultaneously obtained. Interestingly the region within the core contained nonhomogeneous phase maps, also seen in previous studies^14,20^ and raise potential links to described functional inhomogeneities in foveolar vision^63^. Finally, we find that the cortical magnification is large for the topographic foveola (16-25 mm/deg for the central 0.1° and for the central 0.01°, potentially approaching 700 mm/deg, **Fig. 7C,7F**). In addition, if the area of the foveolar core is taken into account this adds another ∼200-300 mm^2^ in area.

**Fig. 8.**
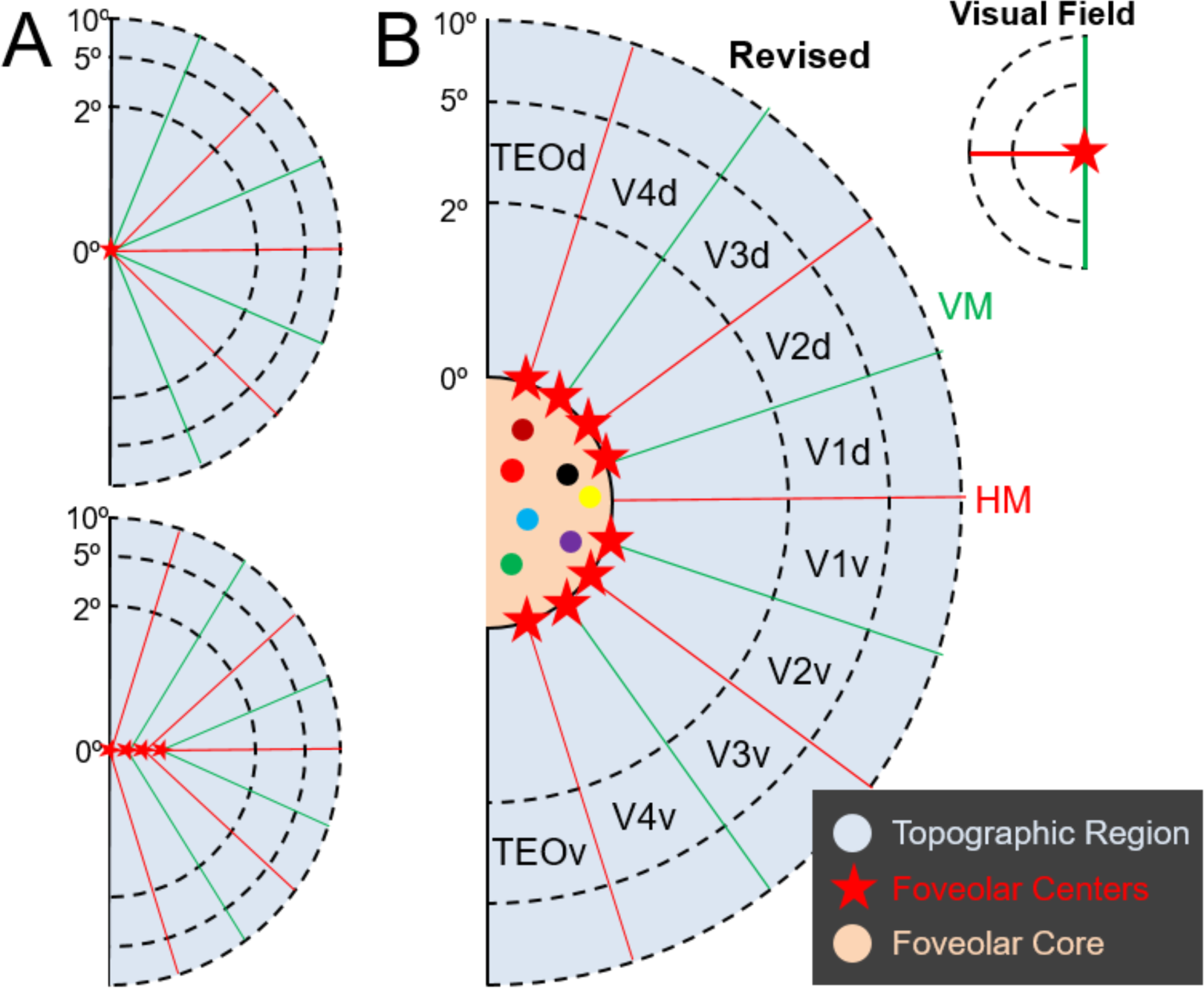
Revised view of visual foveolar representation. **(A)** Previous classical view of topographic representation (light blue). The foveolar confluence (red stars) comprises a single foveolar locus (*Upper*: Zeki 1969^11^) or one locus per area (*Lower*: Schira et al 2009^20^, in humans; Kolster et al 2014^22^, in macaque). **(B)** Our study shows that: (1) the foveolar center is represented 8 times (8 red stars), one at each of the dorsal and ventral representations of V1/V2, V2/V3, V3/V4, V4/TEO. (2) the ‘*foveolar core*’ (orange) is an area within the ring of stars and outside the area of visuotopic representation. (3) there are functional domains (colored dots) within the core that are responsive to large but not small foveolar stimuli (including color, oriented, motion stimuli). Red lines: HM, horizontal meridian. Green lines: VM, vertical meridian.

### Functional domains

Other features also distinguished this central area. The foveolar core was marked by interspersed millimeter scale functional domains. Some domains preferred very high spatial frequencies (11, 15, 18 cyc/deg, higher SF not tested), higher than non-foveolar SFs mapped in primate functional imaging^38,64,65^, but consistent with high spatial frequency preferences of macaque foveolar retinal ganglion cells (15 cyc/deg^66^; 20-40 cyc/deg^67^). These high spatial frequency preference domains could contribute to the high acuity needs of foveolar vision^35,68–70^. Other domains were responsive to color and motion dot stimuli. Notably these domains were not seen with small foveolar stimuli and yet were readily activated by other types of larger stimuli (including high spatial frequency, full field, motion dot stimuli). Unlike other visual areas, the arrangement of these domains is interspersed and does not appear to be similar to that in either V1, V2, V3, or V4. It is possible the populations of functional domains in the core may contribute to non-homogeneous nature of phase encoded maps. While further investigation is needed to understand the function and organization of these domains, our findings suggest marked distinctions between the foveolar core and visuotopic foveolar cortex. *An architectural specialization for foveolar re-representation.* Our findings suggest the foveolar core is a concept distinct from the foveolar confluence. Whereas the confluence refers to a singularity or multiple singularities where multiple cortical areas meet, the core reflects an independent area, with distinct defining features. The foveolar core’s substantial presence (∼200-300 mm^2^, roughly 15-35% the area of V1 in one hemisphere) is an evolutionary statement of its behavioral importance. The architecture of a central zone nestled amongst multiple (8) foveolar representations in each hemisphere encapsulates par excellence the concept of cortical specialization. While re-representation is itself a type of specialization--seen, for example, within areas, in ocular dominance columns in V1, color/shape/disparity stripes in V2, curvature and hue domains in V4, and motion and disparity domains in MT—the foveolar core may exemplify a special coordinator of *multi-areal* re-representation. Although further study is needed, within the core, our findings raise the possibility of that populations of functional domains, distinct from those in areas V1-V4, may be specialized for handling the demands of foveolar vision. Implications of this central architecture for other cortical confluences may also be investigated (cf. MT, parietal confluences).

### A specialization for organizing foveolar behaviors?

We raise a few exciting avenues of investigation into the role of the foveolar core. Two possibilities include (a) *switching between the functional circuits in which* the foveola participates. For example, using the foveola to peruse details of a patterned object (e.g. high spatial acuity and color vision, microsaccades in vision and attention ^5,71^,^72^may recruit substantially different ventral and dorsal pathway or other brainwide circuits than e.g foveal visual updating^73^, or foveally guided fine manual behavior^74,75^. (b) As the two halves of our visual world are represented separately in the two hemispheres and the upper and lower visual fields represented in dorsal and ventral regions of each hemisphere, the foveolar core may be holding a special role in *tying together the different quadrants of our visual and behavioral world*. The unique features of the ‘foveolar core’ call for further investigation into how this cortical specialization participates in primate-specific foveal behaviors^71,76,77^.

## Methods

### Subjects

Animal care and experimental procedures were performed in accordance with the National Institute of Health’s Guide for the Care and Use of Laboratory Animal, and approved by the Institutional Animal Care and Use Committee of Zhejiang University. Two rhesus monkeys (Monkey E: Macaca mulatta 8 –10 kg, male; Monkey J: 4-6kg, female) were used in this study. Monkeys were implanted with MRI-compatible headposts and trained to sit in a natural “sphinx” position inside a plastic box. To motivate monkeys to learn the task, water was provided only during the task performance, 6 days per week. Water was provided *ad libitum* on the 7^th^ day. A daily minimum of water was provided in the chair, and supplemented with a variety of fruit. To ensure that monkeys remain healthy, food intake, urine and feces were monitored daily, and body weight was monitored weekly.

### Behavioral Training

#### Motion Training

The monkeys were trained to sit in a sphinx position in a customized plastic monkey chair^28^. The head was restrained with an implanted headpost secured to the chair. Though the head was restrained, body movements can still produce large motion-related imaging artifacts. We therefore spent some effort on training the monkeys to limit body movements. Our motion training system consisted of 3-dimensional motion sensor (placed on the monkey’s neck) which triggered both negative feedback during body movement (jingling sound, withholding of juice) and positive feedback when keeping still for a set period of time (low frequency sound, juice reward). This training was largely conducted in the mock bore prior to onset of training in the scanner. With training, the time periods for limiting body movement were gradually extended from 1s to 4s; the monkeys learned to adapt and kept still for up to several minutes at a time, a period sufficient for scan acquisitions. Once the monkey learned to keep still in the mock bore, a recorded MRI scanning sound was played so the monkey could adapt to the scanning noise.

#### Fixation Training

Once the monkeys were trained to sit still in the monkey chair, they were placed in a mock scanner, directly facing a liquid crystal display (LCD) screen, which was positioned at 60 cm (the shortest suitable working distance for the eye-tracker) from the monkeys’ eyes. During initial training, they were required to maintain fixation within a 1° window centered on a red dot (0.3°) in the center of the screen. Eye position was monitored at 250 Hz through pupil position and corneal reflection (Eyelink 1000 plus, CR system, Canada). During the training phase, the monkeys were rewarded (fruit juice) for keeping fixation on the red dot within the fixation window for the required time. The fixation duration was gradually extended over time from 0.5s to 3s. After several months of training, both monkeys learned to maintain good fixation lasting 1-3 hours. This resulted in stable activations in response to iso-eccentricity and iso-polar stimuli. The stability of activation was assessed by calculating the RAS (Right, Anterior, Superior) coordinate distance between the center-of-mass across different trials (**Fig. 1D**) in the volume.

#### Scanning Training

Even though the monkey had received training to become accustomed to scanning noise in the mock bore, scanning noise in the actual 7T was still much louder than the recorded ones. To help monkeys adjust gradually, a mild sedative (dexmedetomidine, gradually reduced from 20μg/kg to 8μg/kg) was used over the initial 1-2 weeks. Monkeys were trained on the identical procedures learned in the mock bore. Eye tracking and juice reward methods were identical. Monkeys quickly adjusted and were able to maintain good fixation in scan sessions lasting 1-4 hours. Only data acquired during good fixation (fixation maintained within a 1° radius >85% of the time) was used in this study.

#### Custom RF Coil

For the purpose of enhancing SNR, we built a custom 16-channel surface coil^28^ that greatly enhances the signal-to-noise within a local cortical region. A special acquisition strategy with combined reduced-FOV imaging and high parallel imaging acceleration was used to maximize the spatial encoding capability of the MRI system. This is key to enabling submillimeter resolution functional images (0.6-mm in-plane). While this local transmit coil produces a non-uniform excitation profile, careful positioning of the coil provides a uniform field of foveolar cortex (defined as B1_Max_/ B1_Min_ < 2). A spatial limitation of the coil is that it cannot reliably provide coverage for foveola of both hemispheres; we have therefore focused on a single hemisphere in each animal.

### MRI

#### Anatomical

Scans were performed on a 7T MRI system (Magnetom, Siemens Healthcare, Erlangen, Germany) for all experiments. Whole brain structural images were acquired with single loop coil (RAPID Biomedical GmbH); the monkey was sedated with dexmedetomidine (8-20ug/kg) during these scans. To obtain T1 weighted structural images, we used three-dimensional (3D) magnetization prepared rapid acquisition with gradient echo sequence (MPRAGE) sequence with optimized parameters (TR/TE/TI 2590/2.73/1050ms; matrix size 192×192; FOV 96 mm × 96 mm; slice thickness 0.5 mm; FA 7 degrees; bandwidth 250Hz/px; 3 averages; scan time 24’52’’). Structural images of high quality were generated from averaging across multiple sessions.

#### fMRI

In awake monkeys performing a fixation task, we used a custom designed 16 channel dense multi-array coil, positioned over the occipital portion of the head, to enhance signal-to-noise and provide parallel imaging capability. As a result, required acquisition matrix and phase-encoding steps for a specific spatial resolution was minimized. Enhanced SNR and combined acquisition strategy with reduced FOV and high parallel imaging acceleration rate enabled increase in fMRI spatial resolution over multiple visual cortical areas, including V1, V2, V3, V4, and PIT. Single-shot T2*-weighted GRE-EPI was used for functional scanning with TR/TE 2,000/35ms; matrix size 182×140; FOV 110 mm × 85 mm; spatial resolution 0.6 mm × 0.6 mm × 1 mm; FA 70°; bandwidth 784 Hz/px; echo spacing 1.44ms; parallel imaging acceleration factor R=3 with GRAPPA. Block design was applied with paradigm of 20s blank and 20s stimulus presented (each trial 40s). Each run included 7 trials, making scanning time ∼5’ with auto calibration reference scanning at the beginning. In total 560 TRs (4 runs) were included for one functional mapping. Most of the brain images were acquired in a close to horizontal plane.

Monkey E provided spot imaging data from 32 runs (total 224 trials, 7 trials per run, 4480 TRs) in 4 sessions and provided meridian and iso-eccentricity data from 32 runs (224 trials, 7 trials per run, 4480 TRs) in 6 sessions. Monkey J provided spot imaging data from 21 runs (124 trials, 2280 TRs) in 2 sessions and meridian and iso-eccentricity data from 12 runs (84 trials, 1260 TRs) in 2 sessions.

### Visual Stimuli

#### Stimulus presentation

Stimuli were generated by VPixx software and were projected by a projector (PROPixx, VPixx Technologies, Canada) onto a translucent screen placed in the magnetic bore 60 cm in front of the monkey’s eyes. The projector features a native resolution of 1920 × 1080, and driven with refresh 60 Hz in RGB mode (1440 Hz greyscale) with deterministic timing. Eye movements were monitored by an MR-compatible infrared eye-tracking system (Eyelink 1000 plus). The eye tracker could only track one eye at a time. However, in different sessions, either eye was tracked. Once the animal was trained, no difference in the accuracy of the fixation behavior were observed.

#### Meridian mapping

Our stimuli for direct meridian mapping consisted of a narrow band (either vertical: 20 degrees top to bottom; or horizontal: 30 degrees left to right) with a width of 0.15 degrees; the band contained a red and blue checkerboard grid (0.25 degree × 0.75 degree) alternating at 3 Hz, a stimulus which evokes robust visual response. The fixation point was a 0.2-degree dot presented during visual stimulation and during blank.

#### Iso-eccentricity mapping

Stimuli consisted of paired circular (ring) stimuli at two different eccentricities^50^. By presenting paired well-spaced rings, two distinct iso-eccentricities could be mapped simultaneously without response ambiguity. In addition, in some experiments, maintaining the outer ring constant (e.g. at 4°) across stimuli permitted confirmation of stability and more reliable comparisons of the maps across sessions. Inner ring and outer ring pair stimuli were: Monkey E: 0°/4°, 0.4°/4°, 0.6°/3°, 0.8°/4°, 1.2°/6°, 2°/4°, 1°/5°, 1.4°/7°. Monkey J: 0.8°/4°, 1°/5°, 1.4°/7°). The inner ring consisted of 24 segments of red and blue checkerboard (RGB guns: red 255,0,0; blue 0,0,255; Red CIE coordinate was 252.99, 120.39, 0.46; Blue CIE coordinate was 59.33, 13.53, 330.62) and the outer ring had 32 segments, both alternately flashing at 1Hz. The width of the rings was 0.6 degrees.

#### Foveolar mapping

To directly map the foveal cortical representation, square patches of different sizes (0.4 to 1.0 degrees in diameter) were presented at the center of visual field. For the monkeys to maintain excellent fixation during these scans, a fixation cross (0.3 × 0.1 deg lines) was included at the center of the square patches. The patches alternated between red and blue, a stimulus that is known to be effective for activation of color domains in visual cortex^52^. (RGB guns: red 255,0,0; blue 0,0,255; Red CIE coordinate was 252.99, 120.39, 0.46; Blue CIE coordinate was 59.33, 13.53, 330.62) at 3 Hz.

#### Foveal Functional Domain Mapping

To map different spatial frequency domains, full screen (horizontally extended 30 degrees; vertically extended 20 degrees) of iso-luminant achromatic horizontal gratings with spatial frequencies of 0.2 cycle/degree, 11 cycle/degree, 15 cycle/degree, 18 cycle/degree were presented. The temporal frequency of these gratings was all 1 cycle/sec. A fixation cross (0.3 × 0.1 deg lines) was presented at the center of the square patches to help the monkeys maintain excellent fixation. For motion mapping, 20 random rotating clockwise dots (height: 5 pixels; width: 5 pixels) filling in 2 degree visual field were used. The dots rotated coherently in an angle speed of 45 degrees/sec.

#### Phase-encoding Mapping

To further validate the multiple foveolar loci, very fine iso-eccentricity and iso-polar angle mapping stimuli were used. Continuous expanding/contracting of iso-eccentric rings was presented. Each ring was 0.15° wide and 50 rings in total covered 3°. Each block of expanding rings lasted 50 seconds with 10 seconds of 0.05° constant white dot fixation point before the rings. Continuous clockwise rotating checkerboard wedge (5° wedge) rotated over 50 sec from 90° to −90° in contralateral visual field covering 3° of visual field.

### Data analysis

#### Anatomical image processing and surface reconstruction

To make the functional activations visible along the folding pattern of the cortex, we constructed a cortical surface mesh on an inflated view^78^. Programs implemented in FreeSurfer (v6.0, https://surfer.nmr.mgh.harvard.edu/fswiki/FreeSurferWiki) were customized for monkey cortex at ultra-high field (UHF) MRI. First, the header orientation information was corrected from supine position to sphinx position using *mri_convert*. Due to severe B_1_ inhomogeneity in UHF-fMRI images^79^, the whole brain anatomical images from different sessions were corrected for intensity bias by *N4BiasFieldCorrection*^80^. The bias corrected anatomical images from different sessions were then aligned using rigid body registration by *mri_robust_register* and were averaged for a high-quality anatomical image. The remaining procedures for surface reconstruction were similar to *recon-all* work flow for high resolution data^81^, except that the default human atlas was replaced with D99 monkey atlas^82^.

Similar to de Hollander et al.^83^, the performance of surface reconstruction was carefully inspected and replicated for more than 3 iterations after manual correction for satisfactory surface reconstructions. Crucially, we checked if white matter segmentation and the brain skull strip were correct. Error in white matter segmentation will lead to misposition of white matter/gray matter surface. If the white matter mask extended in to gray matter, then the white matter mask was corrected using *Freeview*. If there was white matter lost, the “control point” was added to give a prior intensity to the intensity bias correction. Error in skull strip will create gray matter outside the brain (e.g., dura). Therefore, dura was manually corrected using *Freeview* where there is a misposition of gray matter/CSF surface.

#### Functional image preprocessing

The preprocessing was performed using AFNI (https://afni.nimh.nih.gov) and FreeSurfer v6.0. DICOM files were transformed to NIfTI data format with AFNI *Dimon* and, same as anatomical image, the header orientation information was corrected from supine position to sphinx position. EPI images were then preprocessed with slice-timing correction, distortion correction with a reverse phase-encoding image^84^ and motion correction. The first frame of the first run in each scan session (4 runs) was used as reference template frame for motion correction using rigid registration with cubic interpolation (using 3dvolreg). Then, the transformations from distortion correction and motion correction were concatenated and applied to the slice timing corrected data only once to reduce the spatial blur induced by interpolation of preprocessing^85^. No extra spatial smoothing was applied on either volume or surface data. For representing the volume data on surface, the spatial transformation from functional image to whole brain anatomical image was calculated by registering the reference frame to whole brain anatomical through a tag-based registration method. Nine to twelve tags distributed at different slices (dorsal, middle, ventral slices) were chosen and confirmed via sagittal view, coronal view and axial view. Applying this spatial transformation, the activation volume resulting from GLM analysis (see next section) was projected to the reconstructed surface at 0.5-mm cortical depth (i.e., middle surface between white matter/gray matter surface and gray matter/CSF surface) using trilinear interpolation by *mri_vol2surf* function in FreeSurfer.

#### GLM analysis

To detect the activated voxels, a general linear model (GLM) analysis was performed on the preprocessed data using *3dDeconvolve* function in AFNI to calculate t-score maps. One regressor representing the experimental design convolved with canonical gamma HRF along with 6 motion regressors to remove residual motion effects were used in the GLM model. Baseline shift was removed with a fifth-order polynomial function during GLM analysis. As we selected only the runs in which fixation performance fell within the 2-degree fixation window for 95% of the run duration, no further regressors based on fixation performance were used during the statistical analysis of the data.

### Foveolar Localization

We predicted that loci of foveolar activation should remain stable across different sizes of foveolar stimuli and across multiple thresholds of significance. To examine this, each monkey was imaged with 3 spot sizes and each image separated into V1/V2, V2/V3, and V3/V4 regions guided by VM and HM shown in **Fig. 2**. Given the simple flashing visual spot stimuli used, activations were strongest for early areas V1 and V2 and weaker for areas V3, V4, and PIT that have preference for more complex stimuli. Thus, each image was then analyzed at thresholds appropriate for each visual area (high, med, and low thresholds producing mm-scale, 5 mm-scale, and cm-scale localization, respectively). The center of foveolar at each area was determined as a. *highest statistical significance b. consistent across different spot sizes c.in the lateralmost regions of visual cortex*.

### Phase-encoding Mapping

To improve SNR, 3 runs (21 trials) of each expanding rings, contracting rings and clockwise rotating wedges were averaged respectively for phase-encoding mapping. The visual field maps were generated by *3dRetinophase* in AFNI with FFT function, which estimate phase based on fundamental frequency of stimuli. The time-courses sampled from ROIs were transformed to frequency domain using Fast Fourier Transform (FFT).

### CMF calculation

Cortical magnification factors were quantified from topographic maps. The linear CMF (deg/mm) was calculated based on brain surface iso-eccentricity maps (12 iso-eccentricity rings) and the points of iso-eccentricity line intersections with the vertical (VM) and horizontal meridian lines. Linear CMF was calculated by measuring the distance (mm) along the cortical surface (using *mris_pmake* from FreeSurfer function) between intersection points of two eccentricities along the meridians divided by the visual distance (deg). Average linear CMF along the polar axis were also calculated based on the measured cortical distance between HM and VM at a given eccentricity and divided by the associated arc length for that eccentricity. Distance between HM and VM at a given eccentricity were generated by adding the distances between adjacent voxels activated along the eccentricity line to follow the surface curvature of the brain. Since the foveola loci were determined and these loci formed a ring-like zone, by connecting these identified foveolar loci, the central areas were outlined and labeled as a mask. The size of this connected area could be calculated by *mris_anatomical_stats* from FreeSurfer function. The single querant 1° areal size in V1 was calculated in the same way by connecting 1° eccentricity line with VM and HM.

### Overlap Calculation

To examine the degree of spatial overlap of functional domains, the Dice coefficient was used to quantify the overlap ratio. Specifically, the number of intersected vertices was divided by the total number of the activated vertices.

### Domain Size Calculation

The area of the domain was calculated by *mris_pmake*. This function overlays the significantly activated voxels onto the surface mesh of the cortex^41^ and determines the number of vertices overlapping with the activation. It then provides the estimated length along the long axis of the domain (see **Extended Data Fig. 8**).

### Intrinsic Optical Imaging

Following craniotomy surgery, the brain was stabilized with agar, and images were obtained through a cover glass. Images of reflectance changecorresponding to local cortical activity were acquired (Imager 3001, Optical Imaging Inc., Germantown, NY) with 630 nm illumination^86^. Signal-to-noise ratio was enhanced by trial averaging (30-50 trials per stimulus condition). Frame sizes were 540×654 pixels representing ∼16×19.4 mm of imaged area. Visual stimuli were presented in blocks. Each block contained all stimulus conditions (e.g., different orientation gratings) and a blank condition, which is a gray screen at the same mean luminance level as grating conditions.

The same gray screen is used for interstimulus intervals (ISI), which were at least 8 s. For each condition, imaging started 0.5 s before the stimulus onset (imaging of the baseline) while the screen remained as a gray blank (the same as in the ISI). Then a visual stimulus was presented for 3.5 s. All stimulus conditions were displayed in a randomized order.

Visual stimuli were created using ViSaGe (Cambridge Research Systems Ltd.) and presented on a 20-in. cathode ray tube monitor (SONY CPD-G520). The stimulus screen was gamma corrected and positioned 57 cm from the eyes. For ocular dominance, orientation maps, full-screen drifting square-wave gratings were used (spatial frequency = 1.5 c/degree, temporal frequency 4 Hz). For color map, responses to redgreen isoluminant sinewave gratings and black-white sinewave gratings (spatial frequency = 1 c/degree, temporal frequency 4 Hz) were compared. Iso-eccentric arcs (0.02° width, radius: 0.01°, 0.1°, 0.2°, 0.3°, 0.4°, 0.5°), and iso-polar bars (0.02° width, angular rotation: 0°, 15°,30°,45°, 60°,75°, 90°) were presented on a uniform gray background at a fresh rate of 8Hz.

For each stimulus condition, we constructed a “single-condition map.” The gray value of each pixel in the “single-condition map” represents the percent change of the light reflectance signal after stimulus onset. These “single-condition maps” were used for calculating difference maps (ocular dominance map: left eye vs. right eye; orientation map: horizontal vs. vertical grating; color map: red/green vs. white/black grating) and vector maps (Color-coded angularity and eccentricity maps).

Color-coded eccentricity and angularity maps (vector summation of conditions) were drawn based on the results of each condition. Each Iso-eccentric arc map/ iso-polar bar map was obtained first. The responses to different eccentricities/angularities at each pixel were then vector-summed to obtain a polar map^87^.

### Mapping foveolar location in anesthetized monkey with optical imaging

The position of the most foveal representation in the image maps was determined by topographic mapping of bars in horizontal and vertical orientation. Following an estimate of the foveal location on the monitor by back projection of the retinal foveolar density visualized through an opthalmoscope, we began by mapping with a sequence of 5 equally spaced bars (each 1° wide) placed at sequential neighboring positions on the screen. Out of the 5 cortical images to these bars, the one which produced the strongest response was identified. This location was then used as a center for testing 5 narrower (0.5°) bars spanning a total of 2.5°. Again, we chose the location of strongest response as the one including the foveolar location and reduced the stimulus size and span by half. This was repeated interatively (1°, 0.5°, 0.2°, 0.1° were tested) to identify the centralmost foveolar location. By imaging sets of horizontal and vertical bars in this fashion, a central foveal location (precision within 0.1°) on the cortex was identified.

**Supplementary Information** is available for this paper.

## Acknowledgments

We thank Hisashi Tanigawa for their help and suggestions on monkey training system. We thank Pinyi Wang for help with scanning and Weihan Li for help with data analysis. This research was supported by National Key Research and Development Program of China (2018YFA0701400) (to A.W.R. and X.Z.); China Brain Project (2021ZD0200401); National Natural Science Foundation of China U1909205 and 31627802 (to A.W.R.), 52277232, 81701774 and 61771423 (to X.Z.), and 32100802 (to J.M.H); Key Research and Development Program of Zhejiang Province 2020C03004 (to A.W.R.); the Fundamental Research Funds for the Central Universities 2019XZZX003-20 (to A.W.R.) and226-2-22-00136 (to X.Z.). China Postdoctoral Science Foundation 2020M681829 (to J.M.H.) and 2020M681866 (to Y.G.); and seed grant of the MOE Frontier Science Center for Brain Science & Brain-machine Integration at Zhejiang University (to A.W.R. and X.Z.)

The authors declare no competing interests.

---

**Extended Data Fig. 1.**
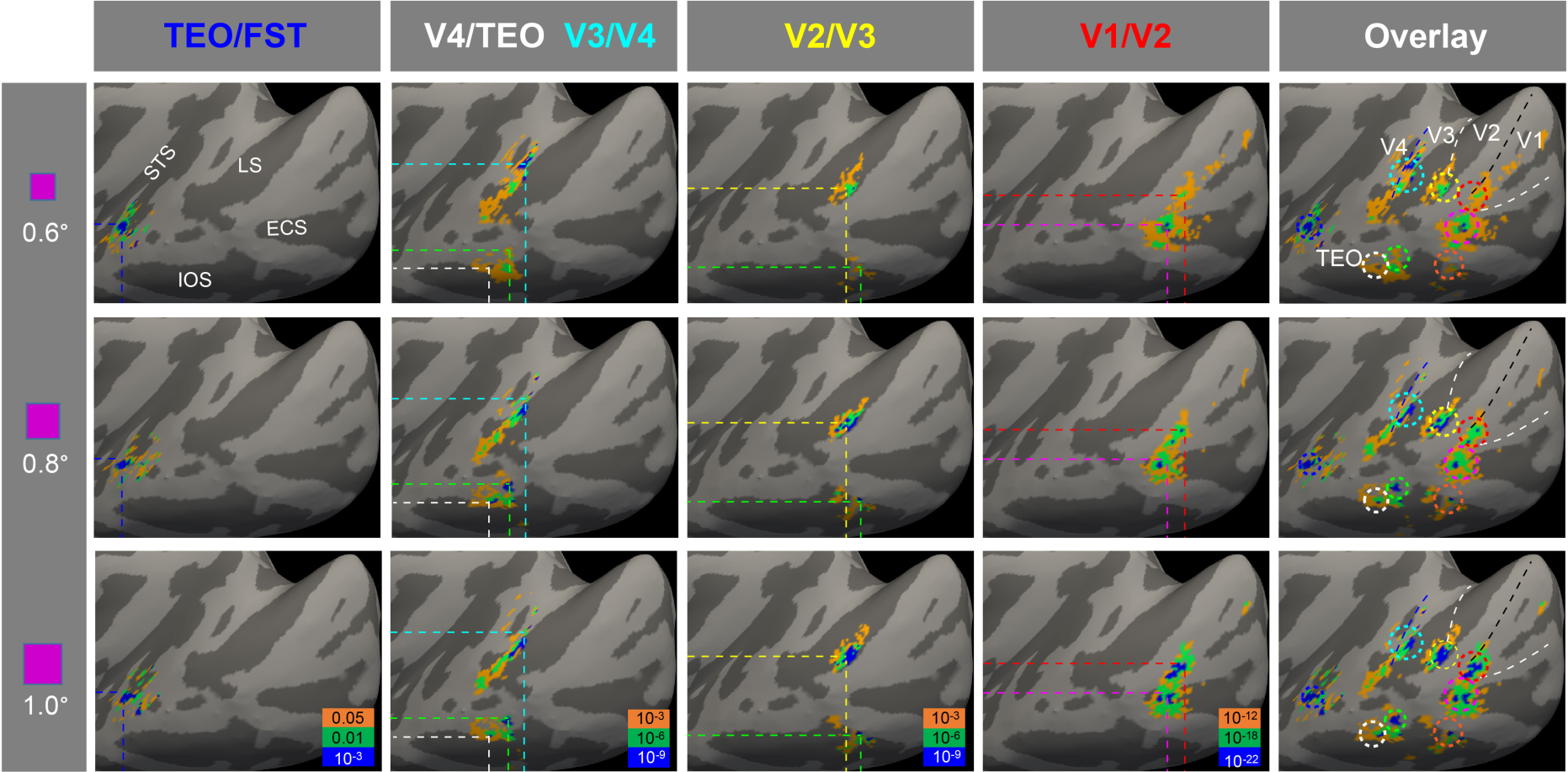
Determining locations of foveolar representation on Monkey J. Shown in each row (V1/V2, V2/V3, V3/V4, V4/TEO, and TEO/FST) are the activation maps to each of the 3 spot stimuli (0.6°, 0.8°, 1.0°). Conventions same as Fig. 3A.

**Extended Data Fig. 2.**
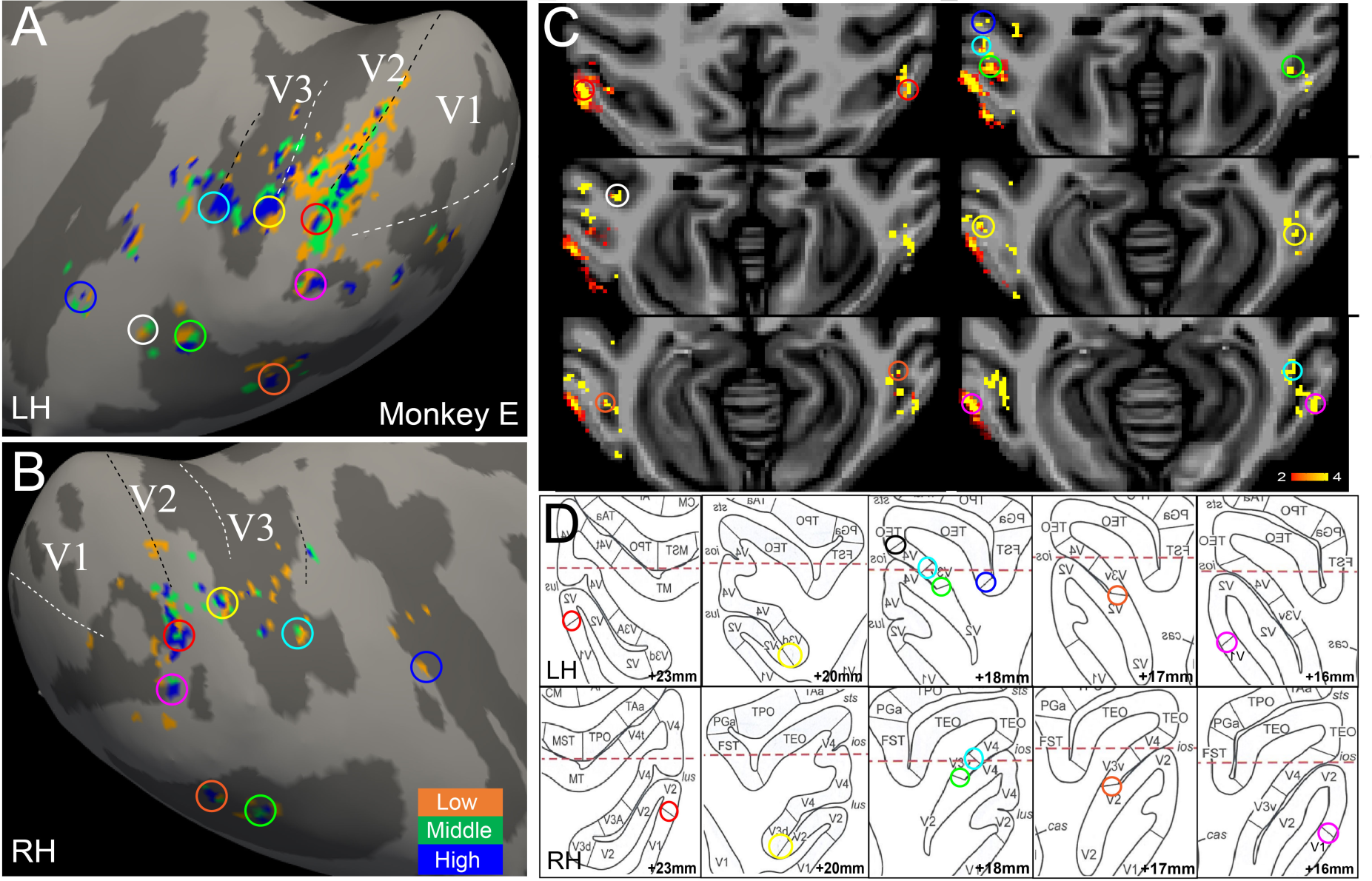
Slice views of bilateral foveolar representation on Monkey E. **(A, B)** Surface view of multiple foveolar representations on the left (A) and right (B) hemispheres, methods same as in Fig. 3. [p-values indicated by color code: orange (lowest), green (middle), and blue (highest)]. (**C**) Slice view of bilateral foveolar activations to spot stimulation of 0.8deg in transverse sections. Color circles refer to the foveolar loci in A&B. (**D**) Corresponding sections to C in D99 atlas. Color circles: foveolar activations corresponded with visual area borders. Numbers: slice number.

**Extended Data Fig. 3.**
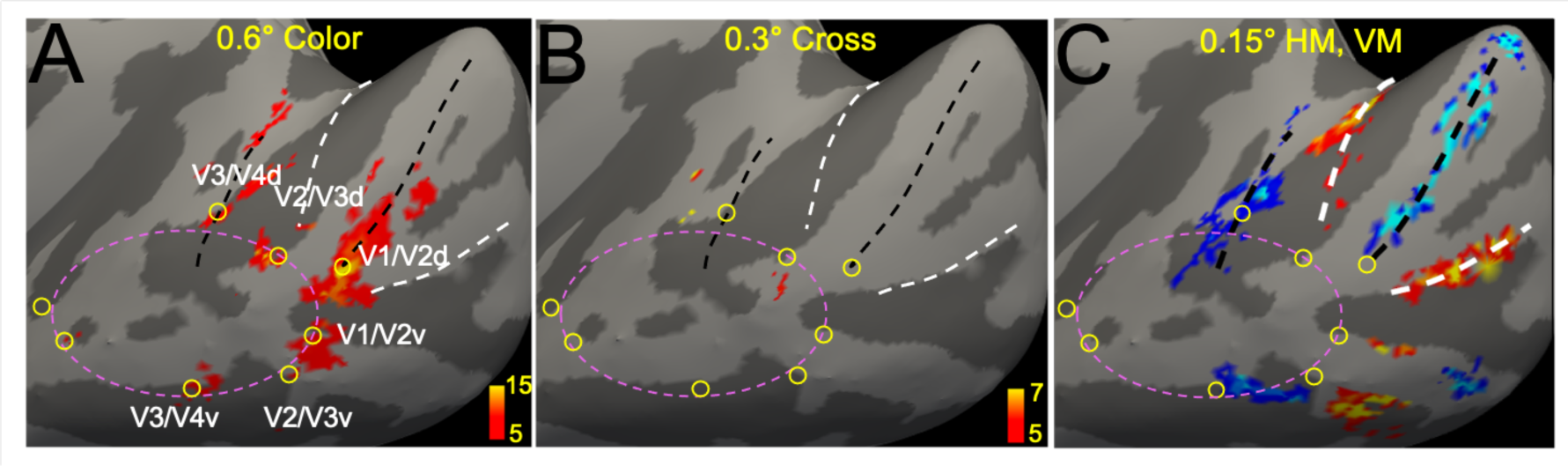
Retinotopic stimuli do not activate foveolar core. Comparison of **(A)** Color alone (0.6° red/blue square), p<10^-5^ evokes multiple foveolar activations (yellow circles) (p<10^-5^, see manuscript **Extended Data Fig. 1** for full map), revealing activation-free foveolar core (dotted pink oval). **(B)** Small fixation cross (lines 0.1° wide, 0.3° long) evokes some weak response at foveolar loci (threshold lowered to p<10^-3^ for visibility), and no activation in core. **(C)** Map of 0.15° wide lines in HM (red) VM (blue) does not invade core area (same as manuscript Fig. 2B).

**Extended Data Fig. 4.**
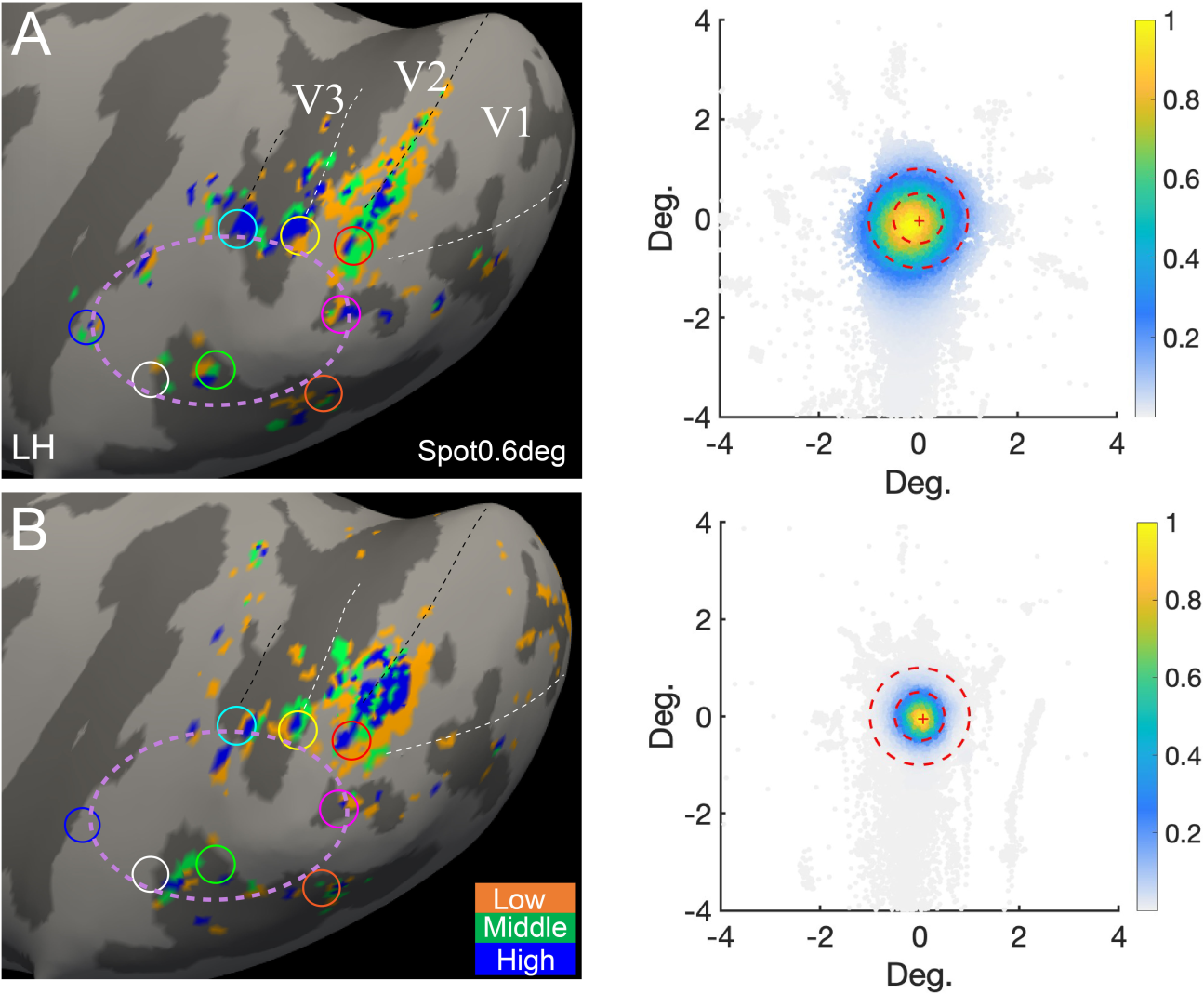
Stability and separability of foveolar loci despite different distributions of eye movements. **(A)** poorer (85% within 1°) and **(B)** better (95% within 1°) eye fixation behavior. Activations on the cortex remain very similar (see circles).

**Extended Data Fig. 5.**
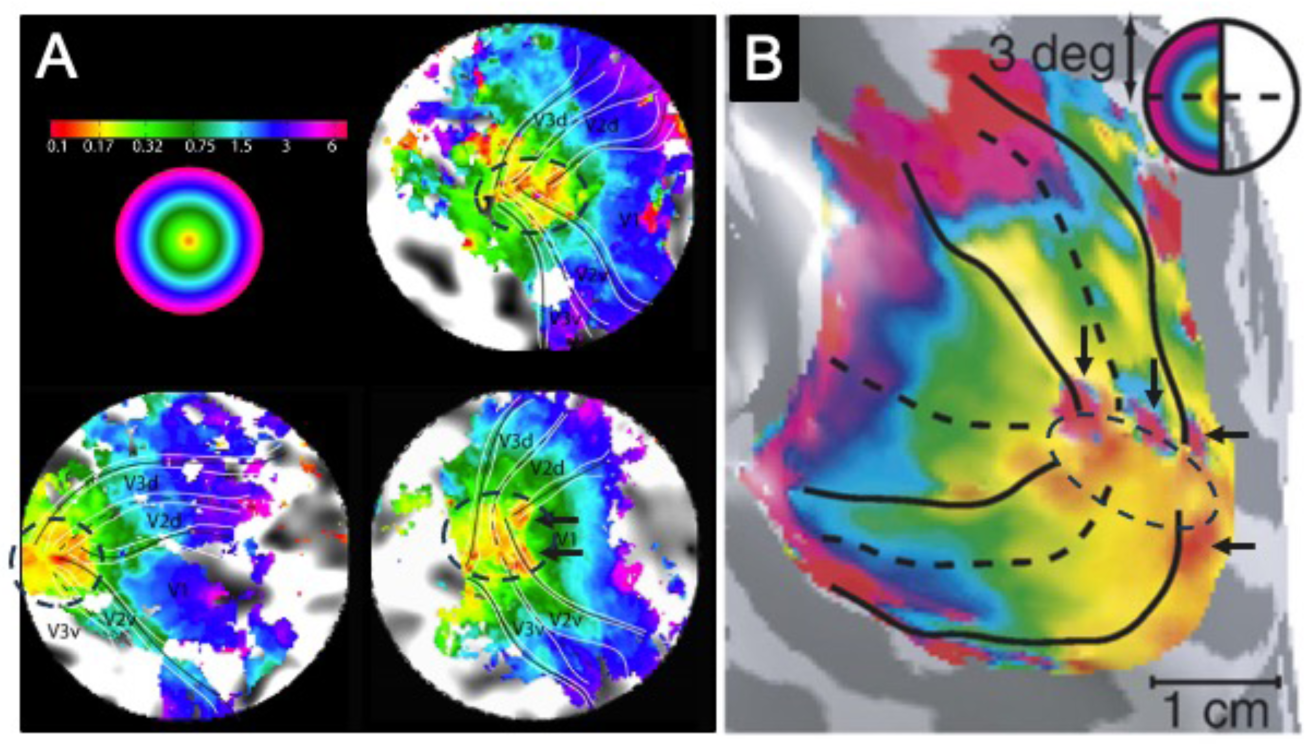
Similarities of human foveolar maps with this study. (A) Phase encoding retinotopic mapping adapted from Schira 2009 Figure 5 reveals nonhomogeneous foveolar region (inside dashed ovals). (B) pRF analysis of retinotopic mapping adapted from Doumulin 2008 Fig. 5. Dashed oval: non-homogenous central area. Note several red foveolar loci at the ends of VM and HM (black arrows) on the dashed ring, an organization similar to what we find.

**Extended Data Fig. 6.**
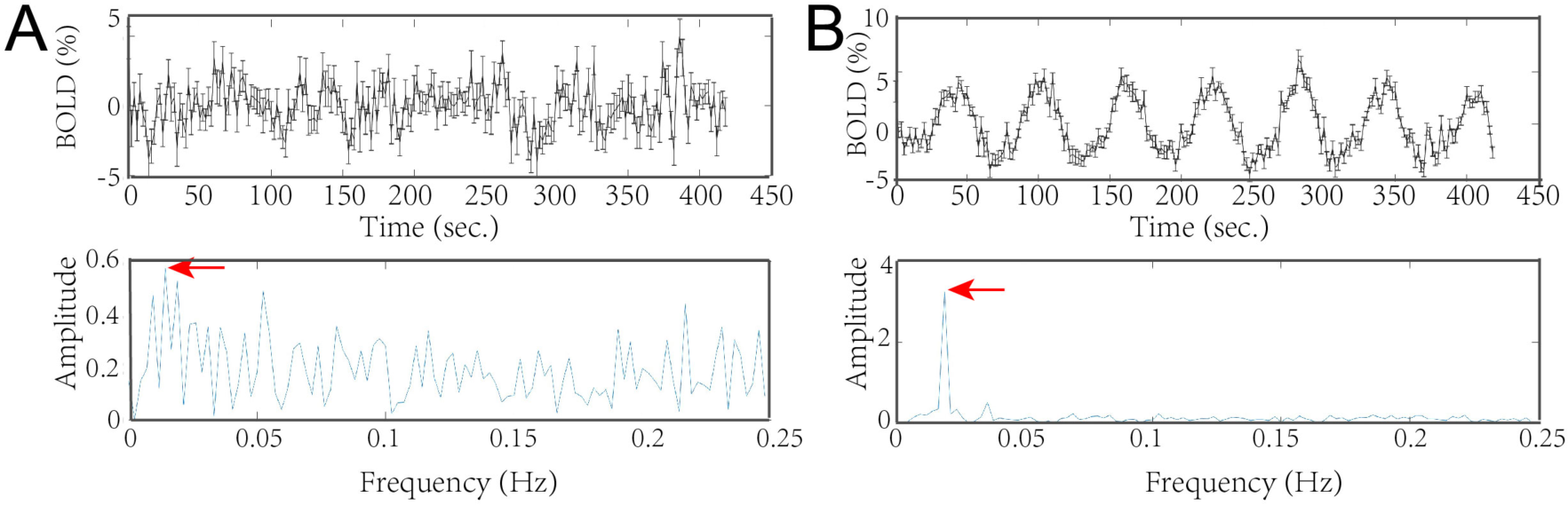
FFT analysis of the time course. Upper panel: Original time-course. Lower pannel: FFT analysis of up time-course. **(A)** Example form ROI #15 (Fig. 4) within the core. (B) Example from ROI #6 (Fig. 4). Red arrow: peak FFT amplitude at stimuli frequency.

**Extended Data Fig. 7.**
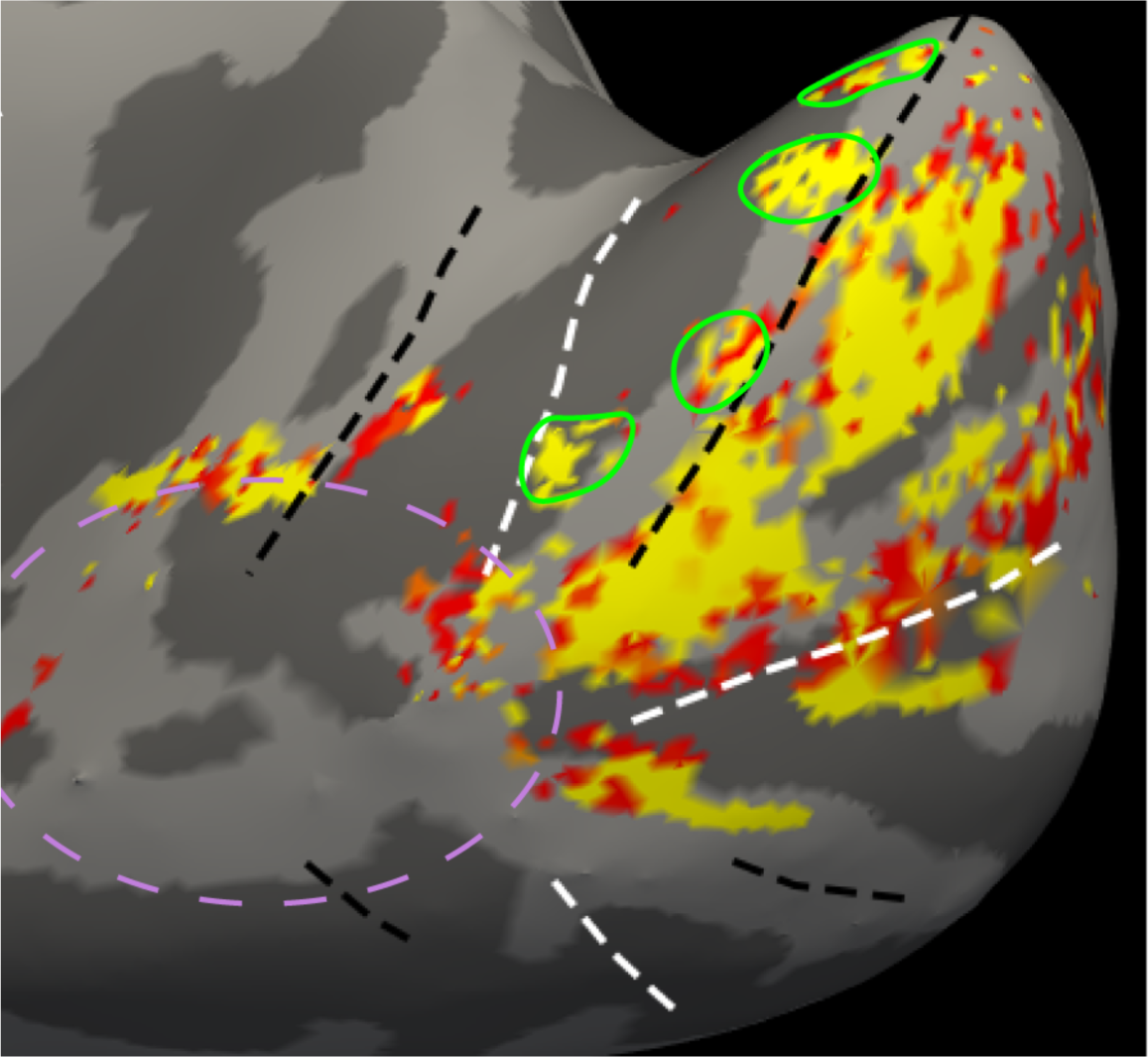
Little activation to standard visual stimuli in the foveolar core. Color stripe mapping in V2. Color grating vs achromatic 1 cyc/deg grating. Green outline: four color stripes in V2.

**Extended Data Fig. 8.**
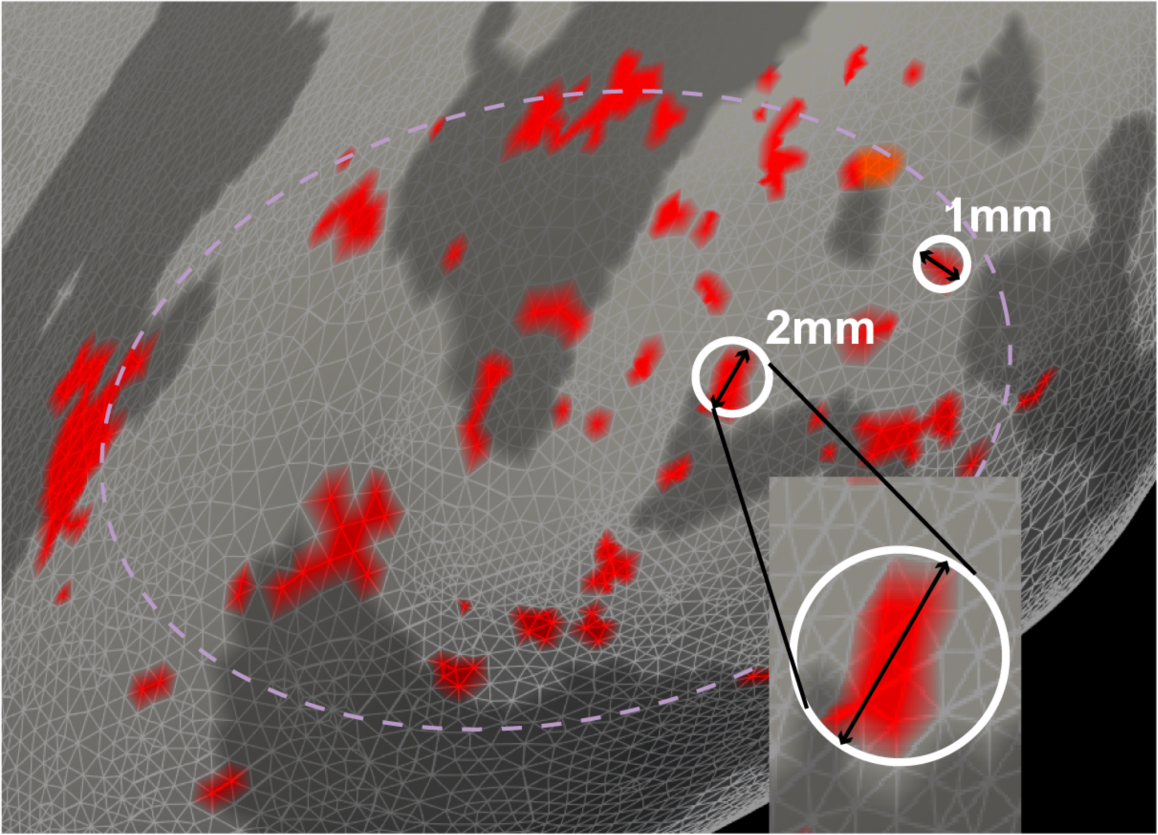
Domain size calculation and distribution. Long axis of domain measured in the surface mesh. Two examples (1 mm and 2 mm size) shown.

**Extended Data Fig. 9.**
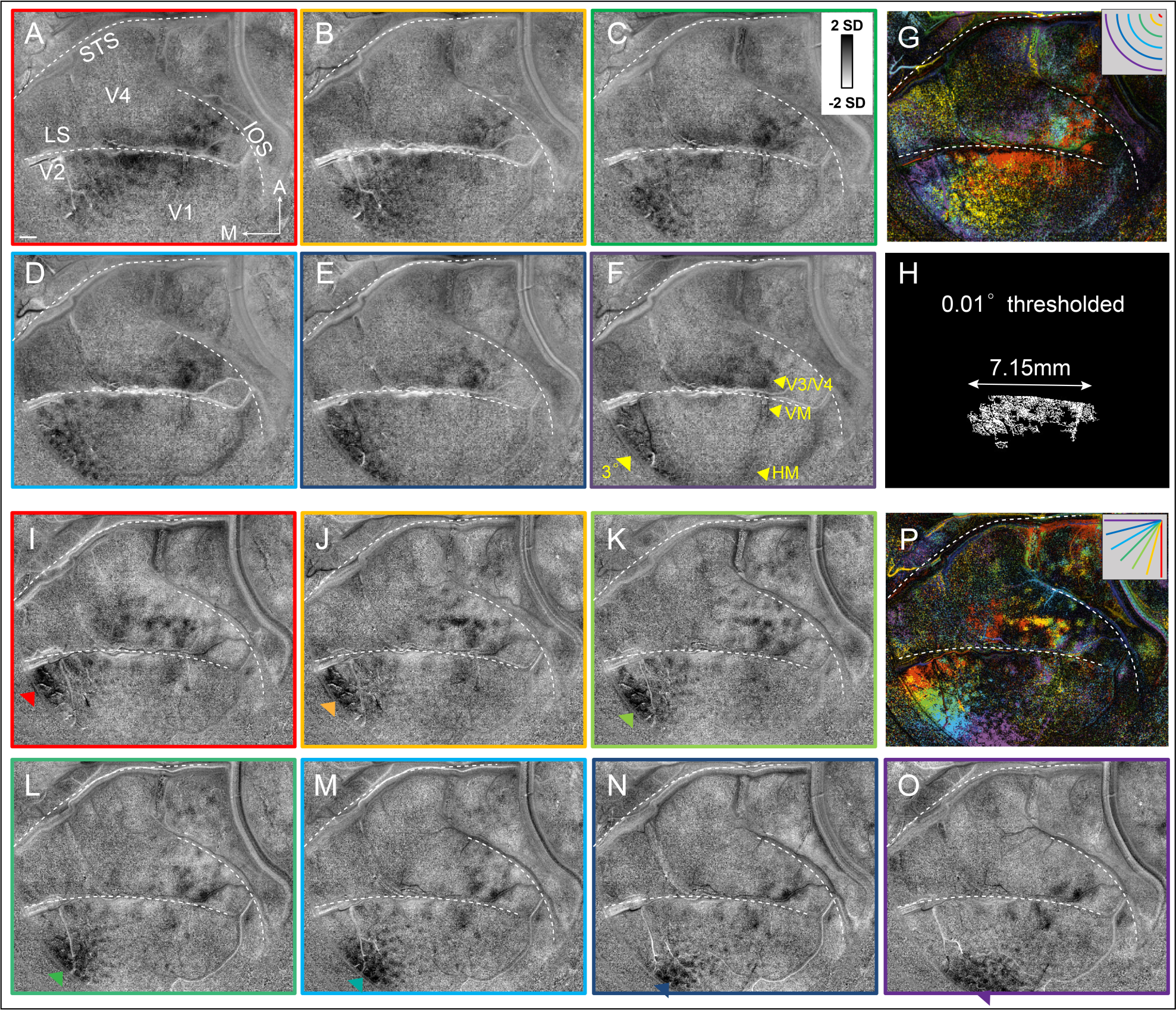
**A-F**. Images (original raw data) of cortical activation to iso-eccentric arcs of radius r in degrees. A: r=0.01°, B: r=0.10°, C: r=0.20°, D: r=0.30°, E: r=0.40°, F: r=0.50°. Scalebar = 1 mm. M: medial, A: anterior, LUS: lunate sulcus, STS: superior temporal sulcus, ISO: inferior occipital sulcus. **G**. Color-coded eccentricity vector map (vector summation of 6 iso-eccentricity responses). **Inset: iso-eccentricity color code**. **H**. **I-O**. Images of cortical activation to isopolar bars in angular rotation Δ in degrees. H: Δ=0°, I: Δ=15°, J: Δ=30°, K: Δ =45 °, L: Δ=60°, M: Δ =75 °, N: Δ=90°. **P**. Color-coded angularity vector map (vector summation of 7 isopolarity responses). (Same case shown in Fig. 4C**-G**.)

## Reference

1. Krubitzer, L. & Kaas, J. The evolution of the neocortex in mammals: how is phenotypic diversity generated? Curr Opin Neurobiol 15, 444–453 (2005).

2. Encyclopedia of neuroscience. (Elsevier, 2009).

3. Roorda, A. & Williams, D. R. The arrangement of the three cone classes in the living human eye. Nature 397, 520–522 (1999).

4. McGregor, J. E. et al. Functional architecture of the foveola revealed in the living primate. PLoS One 13, e0207102 (2018).

5. Poletti, M., Rucci, M. & Carrasco, M. Selective attention within the foveola. Nat Neurosci 20, 1413–1417 (2017).

6. Petersen, C. C. H. The functional organization of the barrel cortex. Neuron 56, 339–355 (2007).

7. Gregory, J. E., Iggo, A., McIntyre, A. K. & Proske, U. Electroreceptors in the platypus. Nature 326, 386–387 (1987).

8. Bichot, N. P., Schall, J. D. & Thompson, K. G. Visual feature selectivity in frontal eye fields induced by experience in mature macaques. Nature 381, 697–699 (1996).

9. McDonald, S. A. & Shillcock, R. C. The implications of foveal splitting for saccade planning in reading. Vision Res 45, 801–820 (2005).

10. Livingstone, M. S. et al. Development of the macaque face-patch system. Nat Commun 8, 14897 (2017).

11. Zeki, S. M. Representation of central visual fields in prestriate cortex of monkey. Brain Research 14, 271–291 (1969).

12. Klink, P. C., Chen, X., Vanduffel, W. & Roelfsema, P. R. Population receptive fields in nonhuman primates from whole-brain fMRI and large-scale neurophysiology in visual cortex. Elife 10, e67304 (2021).

13. Stoll, S., Infanti, E., de Haas, B. & Schwarzkopf, D. S. Pitfalls in post hoc analyses of population receptive field data. Neuroimage 263, 119557 (2022).

14. Dumoulin, S. O. & Wandell, B. A. Population receptive field estimates in human visual cortex. Neuroimage 39, 647–660 (2008).

15. Wandell, B. A., Dumoulin, S. O. & Brewer, A. A. Visual field maps in human cortex. Neuron 56, 366–383 (2007).

16. Wandell, B. A. & Winawer, J. Computational neuroimaging and population receptive fields. Trends Cogn Sci 19, 349–357 (2015).

17. Sereno, M. I. et al. Borders of multiple visual areas in humans revealed by functional magnetic resonance imaging. Science 268, 889–893 (1995).

18. Engel, S. A., Glover, G. H. & Wandell, B. A. Retinotopic organization in human visual cortex and the spatial precision of functional MRI. Cereb Cortex 7, 181–192 (1997).

19. Dougherty, R. F. et al. Visual field representations and locations of visual areas V1/2/3 in human visual cortex. J Vis 3, 586–598 (2003).

20. Schira, M. M., Tyler, C. W., Breakspear, M. & Spehar, B. The foveal confluence in human visual cortex. J Neurosci 29, 9050–9058 (2009).

21. Tootell, R. B. & Hadjikhani, N. Where is ‘dorsal V4’ in human visual cortex? Retinotopic, topographic and functional evidence. Cereb Cortex 11, 298–311 (2001).

22. Kolster, H., Janssens, T., Orban, G. A. & Vanduffel, W. The retinotopic organization of macaque occipitotemporal cortex anterior to V4 and caudoventral to the middle temporal (MT) cluster. J Neurosci 34, 10168–10191 (2014).

23. Conway, B. R. & Tsao, D. Y. Color architecture in alert macaque cortex revealed by FMRI. Cereb Cortex 16, 1604–1613 (2006).

24. Rima, S., Cottereau, B. R., Héjja-Brichard, Y., Trotter, Y. & Durand, J.-B. Wide-field retinotopy reveals a new visuotopic cluster in macaque posterior parietal cortex. Brain Struct Funct 225, 2447–2461 (2020).

25. Engel, S. A. The development and use of phase-encoded functional MRI designs. Neuroimage 62, 1195–1200 (2012).

26. Hesse, J. K. & Tsao, D. Y. Functional modules for visual scene segmentation in macaque visual cortex. Proc Natl Acad Sci U S A 120, e2221122120 (2023).

27. Tootell, R. B. et al. The retinotopy of visual spatial attention. Neuron 21, 1409–1422 (1998).

28. Zhang, X. et al. A 16-Channel Dense Array for In Vivo Animal Cortical MRI/fMRI on 7T Human Scanners. IEEE Trans Biomed Eng 68, 1611–1618 (2021).

29. Roe, A. W. & Ts’o, D. Y. Visual topography in primate V2: multiple representation across functional stripes. J Neurosci 15, 3689–3715 (1995).

30. Xu, X., Collins, C. E., Khaytin, I., Kaas, J. H. & Casagrande, V. A. Unequal representation of cardinal vs. oblique orientations in the middle temporal visual area. Proc. Natl. Acad. Sci. U.S.A. 103, 17490–17495 (2006).

31. Dumoulin, S. O. et al. In vivo evidence of functional and anatomical stripe-based subdivisions in human V2 and V3. Sci Rep 7, 733 (2017).

32. Li, X., Zhu, Q., Janssens, T., Arsenault, J. T. & Vanduffel, W. In Vivo Identification of Thick, Thin, and Pale Stripes of Macaque Area V2 Using Submillimeter Resolution (f)MRI at 3 T. Cereb Cortex 29, 544–560 (2019).

33. Ponce, C. R., Lomber, S. G. & Livingstone, M. S. Posterior Inferotemporal Cortex Cells Use Multiple Input Pathways for Shape Encoding. J Neurosci 37, 5019–5034 (2017).

34. Nasr, S., Polimeni, J. R. & Tootell, R. B. H. Interdigitated Color-and Disparity-Selective Columns within Human Visual Cortical Areas V2 and V3. J Neurosci 36, 1841–1857 (2016).

35. Banks, M. S., Geisler, W. S. & Bennett, P. J. The physical limits of grating visibility. Vision Res 27, 1915–1924 (1987).

36. Dow, B. M., Snyder, A. Z., Vautin, R. G. & Bauer, R. Magnification factor and receptive field size in foveal striate cortex of the monkey. Exp Brain Res 44, 213–228 (1981).

37. Silverman, M. S., Grosof, D. H., De Valois, R. L. & Elfar, S. D. Spatial-frequency organization in primate striate cortex. Proc Natl Acad Sci U S A 86, 711–715 (1989).

38. Nauhaus, I., Nielsen, K. J., Disney, A. A. & Callaway, E. M. Orthogonal micro-organization of orientation and spatial frequency in primate primary visual cortex. Nat Neurosci 15, 1683– 1690 (2012).

39. Aghajari, S., Vinke, L. N. & Ling, S. Population spatial frequency tuning in human early visual cortex. J Neurophysiol 123, 773–785 (2020).

40. Lu, Y. et al. Revealing Detail along the Visual Hierarchy: Neural Clustering Preserves Acuity from V1 to V4. Neuron 98, 417–428.e3 (2018).

41. Winkler, A. M. et al. Measuring and comparing brain cortical surface area and other areal quantities. Neuroimage 61, 1428–1443 (2012).

42. Roe, A. W. et al. Toward a Unified Theory of Visual Area V4. Neuron 74, 12–29 (2012).

43. Hu, J. M., Song, X. M., Wang, Q. & Roe, A. W. Curvature domains in V4 of macaque monkey. eLife 9, e57261 (2020).

44. Conway, B. R., Moeller, S. & Tsao, D. Y. Specialized color modules in macaque extrastriate cortex. Neuron 56, 560–573 (2007).

45. Schira, M. M., Wade, A. R. & Tyler, C. W. Two-dimensional mapping of the central and parafoveal visual field to human visual cortex. J Neurophysiol 97, 4284–4295 (2007).

46. Daniel, P. M. & Whitteridge, D. The representation of the visual field on the cerebral cortex in monkeys. J Physiol 159, 203–221 (1961).

47. Gattass, R., Gross, C. G. & Sandell, J. H. Visual topography of V2 in the macaque. J Comp Neurol 201, 519–539 (1981).

48. Gattass, R., Sousa, A. P. & Gross, C. G. Visuotopic organization and extent of V3 and V4 of the macaque. J Neurosci 8, 1831–1845 (1988).

49. Dow, B. M., Vautin, R. G. & Bauer, R. The mapping of visual space onto foveal striate cortex in the macaque monkey. J Neurosci 5, 890–902 (1985).

50. Tootell, R. B., Switkes, E., Silverman, M. S. & Hamilton, S. L. Functional anatomy of macaque striate cortex. II. Retinotopic organization. J Neurosci 8, 1531–1568 (1988).

51. Hubel, D. H. & Wiesel, T. N. Uniformity of monkey striate cortex: a parallel relationship between field size, scatter, and magnification factor. J Comp Neurol 158, 295–305 (1974).

52. Sincich, L. C., Adams, D. L. & Horton, J. C. Complete flatmounting of the macaque cerebral cortex. Vis Neurosci 20, 663–686 (2003).

53. Kolster, H. et al. Visual field map clusters in macaque extrastriate visual cortex. J Neurosci 29, 7031–7039 (2009).

54. Pfeuffer, J., Merkle, H., Beyerlein, M., Steudel, T. & Logothetis, N. K. Anatomical and functional MR imaging in the macaque monkey using a vertical large-bore 7 Tesla setup. Magn Reson Imaging 22, 1343–1359 (2004).

55. Goense, J. B. M., Ku, S.-P., Merkle, H., Tolias, A. S. & Logothetis, N. K. fMRI of the temporal lobe of the awake monkey at 7 T. NeuroImage 39, 1081–1093 (2008).

56. Newsome, W. T., Maunsell, J. H. & Van Essen, D. C. Ventral posterior visual area of the macaque: visual topography and areal boundaries. J Comp Neurol 252, 139–153 (1986).

57. Maunsell, J. H. & Van Essen, D. C. Topographic organization of the middle temporal visual area in the macaque monkey: representational biases and the relationship to callosal connections and myeloarchitectonic boundaries. J Comp Neurol 266, 535–555 (1987).

58. Gattass, R. et al. Cortical visual areas in monkeys: location, topography, connections, columns, plasticity and cortical dynamics. Philos Trans R Soc Lond B Biol Sci 360, 709–731 (2005).

59. Pinon, M. Area V4 in Cebus monkey: extent and visuotopic organization. Cerebral Cortex 8, 685–701 (1998).

60. Rosa, M. G. P., Piñon, M. C., Gattass, R. & Sousa, A. P. B. Third tier ventral extrastriate cortex in the New World monkey, Cebus apella. Exp Brain Res 132, 287–305 (2000).

61. Rosa, M. G. & Tweedale, R. Visual areas in lateral and ventral extrastriate cortices of the marmoset monkey. J Comp Neurol 422, 621–651 (2000).

62. Rosa, M. G. P. & Tweedale, R. Brain maps, great and small: lessons from comparative studies of primate visual cortical organization. Philos Trans R Soc Lond B Biol Sci 360, 665–691 (2005).

63. Zhang, Y., Shelchkova, N., Ezzo, R. & Poletti, M. Transient perceptual enhancements resulting from selective shifts of exogenous attention in the central fovea. Curr Biol 31, 2698–2703.e2 (2021).

64. Zhang, Y., Schriver, K. E., Hu, J. M. & Roe, A. W. Spatial frequency representation in V2 and V4 of macaque monkey. Elife 12, e81794 (2023).

65. Lu, H. D. & Roe, A. W. Optical imaging of contrast response in Macaque monkey V1 and V2. Cereb Cortex 17, 2675–2695 (2007).

66. Crook, J. M., Lange-Malecki, B., Lee, B. B. & Valberg, A. Visual resolution of macaque retinal ganglion cells. J Physiol 396, 205–224 (1988).

67. Godat, T. et al. In vivo chromatic and spatial tuning of foveolar retinal ganglion cells in Macaca fascicularis. PLoS One 17, e0278261 (2022).

68. Levi, D. M. & Klein, S. A. Spatial localization in normal and amblyopic vision. Vision Res 23, 1005–1017 (1983).

69. Cajar, A., Engbert, R. & Laubrock, J. Spatial frequency processing in the central and peripheral visual field during scene viewing. Vision Res 127, 186–197 (2016).

70. Chung, S. T. L., Legge, G. E. & Tjan, B. S. Spatial-frequency characteristics of letter identification in central and peripheral vision. Vision Res 42, 2137–2152 (2002).

71. Shelchkova, N. & Poletti, M. Modulations of foveal vision associated with microsaccade preparation. Proc Natl Acad Sci U S A 117, 11178–11183 (2020).

72. Poletti, M. & Rucci, M. Active vision: adapting how to look. Curr Biol 23, R718–720 (2013).

73. Ludwig, C. J. H., Davies, J. R. & Eckstein, M. P. Foveal analysis and peripheral selection during active visual sampling. Proc Natl Acad Sci U S A 111, E291–299 (2014).

74. Klein, L. K. et al. Distinct Neural Components of Visually Guided Grasping during Planning and Execution. J Neurosci 43, 8504–8514 (2023).

75. Monaco, S., Gallivan, J. P., Figley, T. D., Singhal, A. & Culham, J. C. Recruitment of Foveal Retinotopic Cortex During Haptic Exploration of Shapes and Actions in the Dark. J Neurosci 37, 11572–11591 (2017).

76. Fan, X., Wang, L., Shao, H., Kersten, D. & He, S. Temporally flexible feedback signal to foveal cortex for peripheral object recognition. Proc. Natl. Acad. Sci. U.S.A. 113, 11627–11632 (2016).

77. Most, S. B., Scholl, B. J., Clifford, E. R. & Simons, D. J. What you see is what you set: sustained inattentional blindness and the capture of awareness. Psychol Rev 112, 217–242 (2005).

78. Dale, A. M., Fischl, B. & Sereno, M. I. Cortical Surface-Based Analysis. NeuroImage 9, 179– 194 (1999).

79. Polimeni, J. R., Renvall, V., Zaretskaya, N. & Fischl, B. Analysis strategies for high-resolution UHF-fMRI data. NeuroImage 168, 296–320 (2018).

80. Tustison, N. J. et al. N4ITK: Improved N3 Bias Correction. IEEE Trans. Med. Imaging 29, 1310–1320 (2010).

81. Zaretskaya, N., Fischl, B., Reuter, M., Renvall, V. & Polimeni, J. R. Advantages of cortical surface reconstruction using submillimeter 7 T MEMPRAGE. NeuroImage 165, 11–26 (2018).

82. Reveley, C. et al. Three-Dimensional Digital Template Atlas of the Macaque Brain. Cereb. Cortex cercor;bhw248v1 (2016) doi:10.1093/cercor/bhw248.

83. de Hollander, G., van der Zwaag, W., Qian, C., Zhang, P. & Knapen, T. Ultra-high field fMRI reveals origins of feedforward and feedback activity within laminae of human ocular dominance columns. NeuroImage 228, 117683 (2021).

84. Andersson, J. L. R., Skare, S. & Ashburner, J. How to correct susceptibility distortions in spin-echo echo-planar images: application to diffusion tensor imaging. Neuroimage 20, 870–888 (2003).

85. Glasser, M. F. et al. The minimal preprocessing pipelines for the Human Connectome Project. NeuroImage 80, 105–124 (2013).

86. Lu, H. D., Chen, G., Tanigawa, H. & Roe, A. W. A motion direction map in macaque V2. Neuron 68, 1002–1013 (2010).

87. Bosking, W. H., Zhang, Y., Schofield, B. & Fitzpatrick, D. Orientation selectivity and the arrangement of horizontal connections in tree shrew striate cortex. J Neurosci 17, 2112–2127 (1997).

